# Distinct mitochondrial DNA single-nucleotide variant signatures in *TOP3A*-deficient cardiomyocytes

**DOI:** 10.64898/2026.08.18.745450

**Authors:** Marianne Gaubert, Halima Alachram, Ipek Ilgin Gönenç, Iñigo Domínguez, Carolin Argyriou, Julia Schmidt, Silke Kaulfuß, Mario Pavez-Giani, Constantin Tilman Schott, Axel Munk, Arne Zibat, Lukas Cyganek, Gökhan Yigit, Bernd Wollnik

**Affiliations:** Institute of Human Genetics, University Medical Center Göttingen, Göttingen, Germany; Cluster of Excellence Multiscale Bioimaging: From Molecular Machines to Networks of Excitable Cells (MBExC), University of Göttingen, Göttingen, Germany; German Center for Child and Adolescent Health (DZKJ), partner site Göttingen, Germany; German Centre for Cardiovascular Research (DZHK), partner site Lower Saxony, Göttingen, Germany; Center for Chromosome Stability, Department of Cellular and Molecular Medicine, University of Copenhagen, Copenhagen, Denmark; Institute for Experimental and Translational Cardiology, University of Giessen, Giessen, Germany; Medical Clinic I, Cardiology-Angiology, University Hospital of Giessen and Marburg (UKGM), Giessen, Germany; Cardio-Pulmonary Institute, Giessen, Germany; Institute for Mathematical Stochastics, University of Göttingen, Göttingen, Germany; Stem Cell Unit, Department of Cardiology and Pneumology, University Medical Center Göttingen, Göttingen, Germany; Fraunhofer Institute for Translational Medicine and Pharmacology, Göttingen, Germany

**Author notes:** Corresponding author: Prof. Bernd Wollnik, MD, Institute of Human Genetics, University Medical Center Göttingen, Heinrich-Düker-Weg 12, 37073 Göttingen, Germany. These authors contributed equally to this work (Marianne Gaubert, Halima Alachram).

## Abstract

The human heart has a continuous and exceptionally high demand for energy, which is met primarily through mitochondrial oxidative phosphorylation. This dependence places the maintenance and integrity of mitochondrial DNA (mtDNA) for proper mitochondrial function at the center of cardiac health, as mtDNA instability has been shown to cause mitochondrial dysfunction, which can ultimately impair cardiac function leading to cardiomyopathy and heart failure. However, the contribution of mtDNA instability to cardiac disease remains poorly understood. A growing number of nuclear-encoded proteins have emerged as essential regulators of mtDNA maintenance, organization and segregation. DNA Topoisomerase 3α (TOP3A) is expressed as two isoforms: one is a nuclear-related isoform involved in nuclear genome maintenance while the other localizes to the mitochondria to preserve mtDNA integrity. Recently, individuals bearing biallelic loss-of-function variants in *TOP3A* manifested phenotypic traits, including cardiomyopathy, typical for mitochondrial dysfunction, supporting a potential mechanistic link between mitochondrial genome instability and *TOP3A*-related cardiac disease. Here, we investigated the effects of TOP3A deficiency on mtDNA maintenance and stability in the context of *TOP3A*-associated cardiomyopathy. We employed isogenic wild-type, TOP3A-knockout and BLM-knockout induced pluripotent stem cells (iPSCs) to generate cardiomyocytes (iPSC-CMs) and established a high-throughput, ultra-deep mtDNA sequencing strategy achieving approximately 500,000× coverage to characterize low-frequency mtDNA mutational patterns. Loss of TOP3A triggered an early burst of low-frequency *de novo* mtDNA single-nucleotide variants during cardiac differentiation, with a striking enrichment within the mitochondrial ribosomal RNA genes, accompanied by a progressive increase in the heteroplasmy of low-frequency mtDNA variants inherited from the common isogenic background. These unique mtDNA signatures were associated with defective mtDNA copy-number expansion and impaired mitochondrial respiration in mature iPSC-CMs. Together, our approach uncovered a previously uncharacterized consequence of TOP3A deficiency and established a link between impaired mtDNA maintenance and mitochondrial dysfunction in *TOP3A*-associated cardiomyopathy.

## Introduction

As a member of the BLM-TOP3A-RMI1-RMI2 (BTRR) complex, DNA Topoisomerase 3α (TOP3A) is instrumental in resolving DNA torsional strain during replication and repair, revealing its critical function in genome maintenance^1^. In addition to this nuclear function, TOP3A also exerts a BTRR-independent role through its mitochondrial isoform, which is required for mitochondrial DNA (mtDNA) replication and maintenance^2^. Notably, TOP3A resolves hemicatenane structures that arise at the origin of H-strand replication (OriH) during mtDNA replication termination^3^. Biallelic loss-of-function variants in *TOP3A* cause a spectrum of disorders ranging from adult-onset mitochondrial disease to an early-onset Bloom syndrome (BSyn)-like disorder^4,5^. BSyn is a rare disorder clinically characterized by primary microcephaly, growth deficiency, and cancer predisposition^6^. Some individuals with BSyn-like disorder also develop atypical BSyn features, including dilated cardiomyopathy and clinical signs of mitochondrial dysfunction^5^. In line with these findings, *in vitro* and *in vivo* studies have linked impaired TOP3A function to mtDNA copy-number depletion, major-arc deletion formation, hemicatenated mtDNA accumulation, and replication-fork stalling^3,7,8^. However, the pathophysiology associated with TOP3A variants remains undefined. In particular, whether loss of TOP3A activity drives the emergence of distinct single-nucleotide variants (SNV) mutational signatures has yet to be investigated. This study explores the mechanisms driving SNV-associated mtDNA maintenance defects, mitochondrial dysfunction and cardiomyopathy associated with TOP3A deficiency. We employed BTRR-complex knockout (KO) induced pluripotent stem cells (iPSCs) to generate cardiomyocytes (iPSC-CMs) as an isogenic human *in vitro* model of *TOP3A*-related cardiac disease. After confirming that TOP3A deficiency impairs mitochondrial respiration, consistent with prior reports^9^, we established a high-throughput ultra-deep mtDNA sequencing strategy achieving ∼500,000× coverage to accurately identify low-frequency mtDNA SNVs and correlate mitochondrial function with mtDNA mutational signatures. This approach enabled us to identify a previously uncharacterized consequence of TOP3A deficiency through the establishment of the first CRISPR-engineered, isogenic human TOP3A-knockout iPSC-CM model of *TOP3A*-associated cardiac disease. Loss of TOP3A resulted in an early burst of low frequency *de novo* mtDNA variants, with a striking enrichment within mitochondrial ribosomal RNA (rRNA) genes. Concurrently, TOP3A deficiency reshaped the heteroplasmy trajectories of pre-existing mtDNA variants inherited from the common isogenic background of the wild-type (WT) parental line. This mitochondrial genomic instability preceded defective mtDNA copy-number expansion and impaired oxidative phosphorylation in mature iPSC-CMs. Collectively, our findings establish TOP3A as a safeguard of both mtDNA sequence integrity and heteroplasmy homeostasis in mitochondria, thus placing mitochondrial genome instability as a potential driver of the cardiac manifestations of *TOP3A*-associated cardiac disease.

## Materials and methods

### iPSC culture and maintenance

Human iPSC lines were previously generated and characterized^10^. The iPSCs were maintained under feeder-free, serum-free conditions using StemMACS™ iPS-Brew XF medium (Miltenyi Biotec) on Matrigel-coated plates (BD Biosciences) in a humidified incubator at 37 °C with 5% CO . Cells were passaged at 90% confluency using 0.5 mM EDTA and cultured with daily medium changes, supplemented with 2 µM Thiazovivin (Merck Millipore) during replating.

### Differentiation of iPSC into iPSC-CMs

Differentiation was initiated at 80-95% iPSC confluency, depending on the cell line. iPSC-CMs were generated through Wnt signaling modulation and subsequent metabolic selection in feeder-free and serum-free culture conditions as previously described^11^. Following selection, pure iPSC-CMs were replated onto Matrigel-coated plates. Cells were harvested or subjected to functional analyses at days 12, 30, 60, and 90 post-differentiation.

### Genomic DNA isolation

iPSC and iPSC-CM pellets were collected, snap-frozen in liquid nitrogen and stored at -80 °C. Genomic DNA was extracted using the DNeasy Blood & Tissue Kit (QIAGEN) according to the manufacturer’s protocol, with the exception that DNA was eluted in 100 µL AE buffer. DNA concentrations were determined using a NanoDrop™ One Spectrophotometer (Thermo Fisher Scientific).

### Mitochondrial DNA-enriched library preparation and ultra-deep mtDNA sequencing

Mitochondrial DNA-enriched sequencing libraries were prepared from genomic DNA using the Agilent SureSelect XT HS target enrichment system. Genomic DNA was quantified by Qubit fluorometry, and approximately 100 ng was used as input. DNA was first enzymatically fragmented on a thermocycler (Bio-Rad C1000) using the SureSelect XT HS Enzymatic Fragmentation Kit (37 °C for 15 min, then 65 °C for 5 min). The fragmented DNA was loaded onto the Magnis NGS Prep System, which performed the remaining library preparation as an automated protocol comprising end repair and dA-tailing, ligation of adapters carrying a sample index and a unique molecular identifier (UMI), pre-capture PCR amplification (8 cycles), hybridization capture of mitochondrial targets with a custom SureSelect biotinylated-probe panel, streptavidin-bead capture and washing to remove off-target fragments, and post-capture PCR amplification (12 cycles). Library concentration and fragment size distribution were assessed on a 4200 TapeStation (Agilent Technologies). Libraries were sequenced on an Illumina NovaSeq platform targeting ultra-deep coverage (∼500,000× mean raw depth per sample) to enable the detection of low-frequency mtDNA variants.

### Mitochondrial DNA variant calling pipeline

The complete sequencing-data preprocessing and variant-calling workflow is summarized in supplementary Figure 1A and B. Raw sequencing reads were assessed for quality with FastQC (v0.11.5^12^). Unique molecular identifiers (UMIs) were extracted and sequencing adapters trimmed using AGeNT (v3.1.1, Agilent Technologies). Reads were aligned with BWA-MEM (v0.7.17-r1188^13^) to both the standard revised Cambridge Reference Sequence (rCRS, NC_012920.1) and a “shifted” reference in which the coordinate origin is moved by ∼8 kb, the latter enabling accurate variant detection across the artificial linearization breakpoint of the circular mitochondrial genome. UMI-based consensus reads were generated separately for each alignment using AGeNT CReaK (v3.1.1). Sequencing was performed targeting a mean depth of 500,000× per sample; following UMI-based consensus generation, the achieved mean genome-wide consensus depth across all analyzed samples was ∼600,550× (SD ≈ 76,710×; range 516,900-809,400×; n = 15 samples), with no systematic difference in depth between genotypes (Supplementary Figure 2A and B). Indel quality scores were annotated using LoFreq’s indelqual module (--dindel mode) to improve indel-calling sensitivity. Variants were called independently against each reference using LoFreq (v2.1.5^14^), which is a sequence-quality-aware variant caller validated for the detection of low-frequency heteroplasmic variants. Rather than a fixed minimum variant-allele-frequency (VAF) cutoff, calls were determined by LoFreq’s internal statistical significance model, which tests read-level mismatches against the local sequencing-error background (default significance threshold, sig = 0.01), combined with base/mapping quality thresholds (-q 20, -Q 20, -m 20) and LoFreq’s default post-call filters (minimum coverage, strand-bias). Calls from the shifted alignment were lifted over to rCRS coordinates using Picard LiftoverVcf (v3.4.0), restricted to the region spanning the linearization breakpoint (chrM:16025-16569,1-575), and merged with calls from the direct rCRS alignment restricted to the complementary, non-overlapping region (chrM:576-16024) using bcftools concat (v1.5^15^), yielding a single non-redundant call set per sample. Variant call quality, haplogroup assignment, and potential sample contamination were evaluated using bcftools stats (v1.5), Haplogrep3 (v3.2.2^16^), and haplocheck (v1.3.3^17^). Potential interference from nuclear mitochondrial DNA sequences (NUMTs) was assessed directly by competitive alignment of reads to the full hg38 assembly (Supplementary Figure 2D and E). Across all samples, a mean of ∼97% of reads mapped with high confidence to chrM, with the residual ∼3% mapping to the nuclear genome. Of these nuclear-mapped reads, 46-62% clustered at a single locus, chr1:600,000-700,000, corresponding to a known chr1 mega-NUMT sharing 98.5% sequence identity with mtDNA positions m.3911-9755 (encompassing *MT-ND1/2*, *MT-CO1/2/3*, *MT-ATP6/8*). The maximum false allele fraction attributable to NUMT mismapping ranged from 0.37% to 0.51% across samples (Supplementary Figure 2D) and did not vary systematically with genotype or time point, indicating that NUMT-derived signals constitute a fixed technical background across all samples. Variants were annotated for genomic context and predicted functional consequence using Ensembl VEP (v113.0, cache 113_GRCh38 on GRCh38.p14, Ensembl 113) and MitImpact (v3.1.3).

### Downstream mtDNA variant classification and analysis

First, variants with allele frequency below 0.03% (0.0003) were excluded, a conservative threshold set above the empirical detection limits reported for low-frequency variant calling using UMI-based error correction (as low as 0.025% VAF^18^), ensuring that retained variants are unlikely to reflect residual sequencing or PCR noise. Second, following the gnomAD v3.1 mtDNA pipeline^19^, variants overlapping six positions known to produce systematic technical artifacts were excluded: positions 301, 302, 310, and 316 (within the homopolymer tract at chrM:300-317), position 3107 (the rCRS placeholder “N”), and position 16182 (within the homopolymer tract at chrM:16180-16193); these artifacts arise from Illumina sequencing errors in homopolymer stretches and from misalignment to the N base in the reference. After filtering, the mtDNA mutation load was defined as the number of distinct variant sites called in a genotype, so each reported value is an independent property of a single library. Pre-existing variants were defined, within each genotype independently, as bona fide LoFreq calls present at the iPSC baseline (day 0); pre-existing status did not require concordance across other isogenic lines or replicate clones. These baseline variants were then tracked longitudinally, failure to call a pre-existing site at a later time point did not alter its baseline classification. *De novo* variants were defined by three criteria: (i) no LoFreq call in any isogenic line at day 0, (ii) no LoFreq call in all other isogenic cell lines at any analyzed time point, and (iii) continued detection at every subsequent analyzed time point after first appearance.

### Heteroplasmy trajectory and rate-of-change analysis

Heteroplasmy refers to the fraction of mtDNA copies in a cell that carry a given variant, measured here as allele frequency and reported as a percentage. For longitudinal between-genotype comparisons of heteroplasmy dynamics, we restricted the analysis to variants detected at AF ≥ 0.03% across days 0, 12, 30, 60, and 90 in each genotype. This ensures that each variant contributes five data points to the rate-of-change regression and that paired between-genotype tests compare identical genomic positions across conditions. It yielded 545 variants tracked in all three genotypes at all time points. Variants with a maximum AF ≥ 90% in any condition were additionally excluded, as near-homoplasmic variants show minimal variance in allele frequency over time. For each variant and each genotype, the heteroplasmy rate of change was estimated as the slope of an ordinary least-squares linear regression of allele frequency against time (in days), reported in units of % per day. All 545 variants had detected allele frequencies at all five time points in all three genotypes, providing five data points per regression. Rates were compared between genotypes by Wilcoxon signed-rank test on matched genomic positions for each of the three pairwise comparisons.

The association of rate of change with iPSC baseline heteroplasmy (allele frequency at day 0) was assessed by Spearman rank correlation. Rate distributions were also examined descriptively across mitochondrial gene categories (Complex I, III, IV, V, rRNA, D-loop) and by substitution class (transition vs. transversion).

### PacBio HiFi whole-genome sequencing and mtDNA copy-number estimation

High-molecular-weight genomic DNA was isolated from iPSC-CM pellets using the Monarch HMW DNA Extraction Kit for Cells & Blood (New England Biolabs; T3050) according to the manufacturer’s low-input protocol for cultured cells. PacBio HiFi whole-genome library preparation and sequencing were performed as previously described^20^.

Sequencing data were processed using the Lucid Genome Suite (Lucid Genomics GmbH). HiFi reads were aligned to the GRCh38 no-alt analysis-set reference genome using minimap2 (v2.24^21^) with the map-hifi preset. Per-position sequencing depth was calculated using samtools depth with the parameters - a and -Q 20. The mean sequencing depth across the mitochondrial genome was normalized to the mean depth derived from the diploid nuclear reference loci *ALB*, *B2M*, *RPP30* and *TBP*. The resulting ratio was multiplied by two to estimate the number of mtDNA copies per diploid cell.

### Quantitative PCR-based mtDNA copy number assay

Genomic DNA was diluted to generate three input amounts per sample: 10 ng, 5 ng, and 2.5 ng DNA/sample. Quantitative PCR (qPCR) reactions were prepared in a final volume of 10 µL, comprising 1 µL of forward/reverse primer mix at 10 µM each, 5 µL of QuantiNova™ SYBR® Green PCR Master Mix (2×; QIAGEN), 0.2 µL of ROX Reference Dye (QIAGEN), the appropriate volume of genomic DNA template, and nuclease-free water to volume. Relative mitochondrial DNA copy number was determined by normalizing the Ct value of *MT-ND1* to nuclear reference loci *ALB* and *F8*, on three independent differentiations per genotype and time point (n = 3). Genotypes were compared at each time point by two-way ANOVA with Šídák’s multiple comparisons test. Mean differences, 95% confidence intervals and multiplicity-adjusted *p*-values are reported. Quantification and statistical analysis were performed using GraphPad Prism (v11).

### SDS-PAGE and Western blot analysis

Protein lysates from iPSCs were mixed with LDS Sample Buffer and Sample Reducing Agent (Invitrogen), adjusted to 15 µg protein per sample and denatured at 95 °C for 5 min. Proteins were separated using Mini-PROTEAN TGX Stain-Free gels (Bio-Rad) and transferred to PVDF membranes using the Trans-Blot Turbo Transfer System (Bio-Rad). Membranes were blocked for 1 hour in 5% milk powder in 0.1% Tween 20 in Tris-buffered saline (TBS-T) and incubated overnight at 4 °C with antibodies against TOP3A (1:500; Proteintech, 14525-1-AP) and GAPDH (1:1,000; Cell Signaling Technology, 2118). Following incubation with the appropriate HRP-conjugated secondary antibodies, protein bands were detected using WesternBright ECL substrate (Advansta) and visualized with the ChemiDoc Touch Imaging System (Bio-Rad).

### Mitochondrial respiration assay

iPSC-CMs were seeded in XF24 Seahorse assay plates at a density of 100,000 cells per well approximately 2-3 weeks before day 60 of differentiation. At day 60, mitochondrial respiration was assessed using a Seahorse mitochondrial stress test. Cells were incubated in XF assay medium (102365-100, Agilent) supplemented with 10 mM glucose and 1 mM sodium pyruvate at 37 °C without CO prior to the assay. Following baseline oxygen consumption rate (OCR) measurements, oligomycin A (1 μM; 75351, Sigma-Aldrich), carbonyl cyanide-4-(trifluoromethoxy)phenylhydrazone (FCCP, 0.5 μM; C2920, Sigma-Aldrich), and rotenone/antimycin A (1 μM each; R8875 and A8674, Sigma-Aldrich) were sequentially injected. Cellular respiration was measured using a Seahorse XF24-3 Analyzer, and OCR was normalized to total protein content per well. Quantification and statistical analysis were performed using GraphPad Prism (v11). A one-way ANOVA with Tukey’s HSD post-hoc test was performed on the measurements from three independent iPSC-CM differentiations (n = 3 per genotype). Individual wells constituting technical replicates were displayed as boxplots (WT, n = 14; TOP3A KO, n = 13; BLM KO, n = 13) giving the median and interquartile range, with kinetic traces shown as mean ± SEM.

## Results

### Loss of TOP3A results in mitochondrial dysfunction in mature iPSC-CMs

To model *TOP3A*-associated cardiac disease in an isogenic human system, the parental WT line and previously generated CRISPR/Cas9-edited TOP3A knockout (TOP3A KO; *TOP3A*^c.2473ins/c.2475del7^) and BLM knockout (BLM KO; *BLM*^c.715del2/c.716del^) iPSC lines^10^ were differentiated into iPSC-CMs in three independent experiments (n = 3). Following replating and maturation of the iPSC-CMs until day 60, a stage at which they become increasingly reliant on oxidative phosphorylation (OXPHOS), mitochondrial respiration was assessed using the Seahorse mitochondrial stress test (Figure 1A). BLM KO iPSC-CMs served as a BTRR-complex specificity control, which has an important function on nuclear DNA, but has no established role in mtDNA replication or maintenance. The normalized OCR measurements demonstrated that TOP3A deficiency resulted in impaired OXPHOS function in TOP3A KO iPSC-CMs (Figure 1B). Specifically, basal respiration was significantly reduced compared to WT (p = 0.0319, Figure 1C), alongside ATP-linked respiration (p = 0.0264, Figure 1D). While not reaching statistical significance, maximal respiratory capacity also showed a downward trend in TOP3A KO iPSC-CMs (p = 0.2507, Figure 1E). BLM KO iPSC-CMs remained comparable to WT (p = 0.3021, p = 0.1809, p = 0.8784). Loss of TOP3A therefore compromises mitochondrial respiratory function in iPSC-CMs, thus validating the accuracy of our model to study *TOP3A*-related cardiomyopathy.

**Figure 1.**
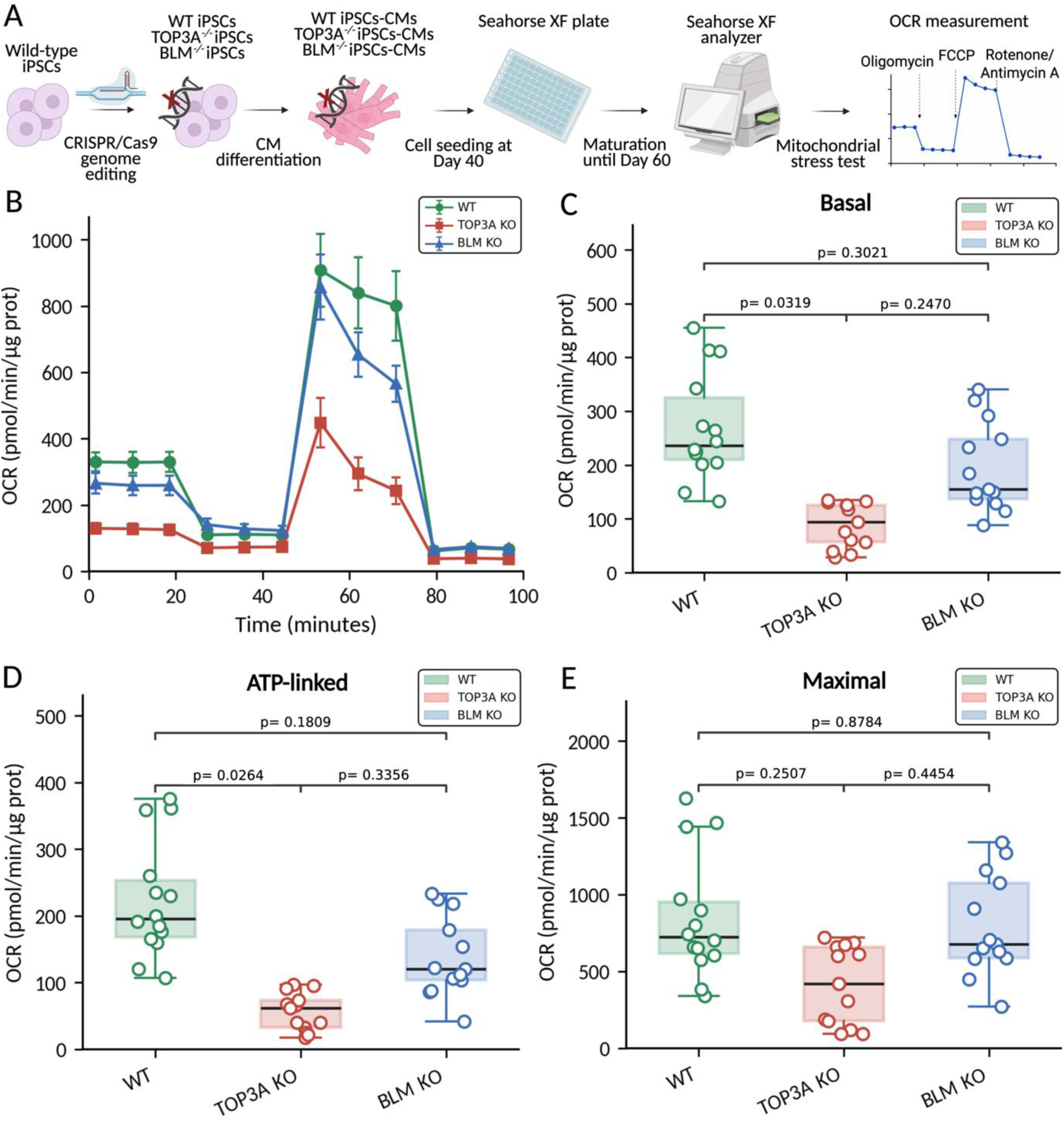
Loss of TOP3A impairs mitochondrial respiration in mature iPSC-CMs. **(A)** Experimental workflow. WT, TOP3A KO, and BLM KO iPSCs generated by CRISPR/Cas9 genome editing were differentiated into iPSC-CMs, seeded into Seahorse XF24 plates at day 40, matured to day 60, and analyzed using the Seahorse XF mitochondrial stress test following sequential injection of oligomycin, FCCP, and rotenone/antimycin A. **(B)** Oxygen consumption rate (OCR; pmol/min/µg protein) as a function of time, normalized to total protein per well, across n = 3 independent differentiations. Points and error bars represent mean ± SEM. WT, TOP3A KO, and BLM KO are shown in green, red, and blue, respectively. **(C-E)** Boxplots showing basal respiration **(C)**, ATP-linked respiration **(D)**, and maximal respiration **(E)**. Boxes indicate the median and interquartile range, and open circles represent individual technical replicate wells (WT, n = 14; TOP3A KO, n = 13; BLM KO, n = 13) pooled across differentiations. Genotype comparisons were performed using one-way ANOVA followed by Tukey’s multiple-comparisons test.

### Mitochondrial DNA mutation load increases upon differentiation in TOP3A KO iPSC-CMs

Given the role of TOP3A in mtDNA maintenance, we next investigated whether mitochondrial dysfunction in TOP3A KO iPSC-CMs coincided with a distinct mtDNA mutational signature. We isolated genomic DNA from WT, TOP3A KO and BLM KO cells at day 0 (iPSCs) and at days 12, 30, 60, and 90 after cardiac differentiation into iPSC-CMs as described^11^. Following library preparation, the enriched mtDNA fractions were subjected to ultra-deep mtDNA sequencing enabling the detection of low frequency mtDNA variants (Figure 2A). To ensure robust variant detection while minimizing amplification and sequencing artifacts, reads underwent stringent quality assessment, UMI-based consensus generation, and sequence-quality-aware variant calling. The achieved mean UMI-consensus depth was approximately 600,550× across all samples, with no systematic differences between genotypes; the mean TOP3A KO/WT coverage ratio was 1.015 ± 0.151 and did not differ from 1 by either one-sample t-test or Wilcoxon signed-rank test (Supplementary Figure 2A and B). In contrast, BAM-level allele frequencies at nine representative variant positions were consistently higher in TOP3A KO, with a mean TOP3A KO/WT allele-frequency ratio of 4.42 ± 2.70, which differed significantly from 1 by both one-sample t-test and Wilcoxon signed-rank test (Supplementary Figure 2B). At this depth, the 0.03% allele-frequency threshold applied in downstream analyses corresponds to approximately 180 supporting consensus reads at mean coverage. Competitive alignment to the complete hg38 reference genome showed that 96-97% of reads mapped to chrM, whereas only ∼3% mapped to the nuclear genome. Among these nuclear-mapped reads, the maximum allele frequency potentially attributable to NUMT misalignment ranged from 0.37% to 0.51% and remained comparable across genotypes and time points (Supplementary Figure 2D and E). At the iPSC level, mtDNA mutation load was comparable across the three isogenic lines, consistent with a largely shared mitochondrial genetic background (Figure 2B; Supplementary Figure 2C). However, after differentiating into iPSC-CMs, the mutation load increased exclusively in TOP3A KO, exceeding 2,500 variants (confident LoFreq variant calls) by day 12 and reaching a maximum of 2,714 variants at day 30, before remaining persistently elevated through days 60 and 90. In contrast, the mutation load progressively declined in WT and BLM KO to minima at day 60 (775 and 588 variants, respectively), followed by a partial increase at day 90. These differences could not be attributed to sequencing depth or NUMT misalignment, as illustrated in Supplementary Figure 2. Across WT, TOP3A KO, and BLM KO genotypes, SNVs constituted the predominant variant class, accounting for 97.05% of all variants detected across genotypes and time points (Supplementary Figure 3A). The remaining 2.95% comprised short indels. Deletions were slightly more frequent than insertions, with TOP3A KO showing the highest numbers of both insertions (41 versus 33 in WT and 28 in BLM KO) and deletions (53 versus 41 and 38, respectively). Indel lengths were comparable across genotypes, averaging approximately 1.5 bp for deletions and 1.3 bp for insertions (Supplementary Figure 3B). Although most indels were shared across genotypes, 23 were detected exclusively in TOP3A KO, of which 20 localized to the D-loop or the mitochondrial rRNA genes *MT-RNR1* (12S) and *MT-RNR2* (16S) (Supplementary Figure 3C). Given their comparatively low abundance, subsequent analyses focused primarily on SNVs, enabling a detailed characterization of the SNV landscape associated with TOP3A deficiency. Transitions predominated over transversions in both WT and TOP3A KO, accounting for 87.4% and 82.0% of SNV calls, respectively, across days 12 to 90 (Supplementary Figure 3D). Both transitions occurred preferentially on the H strand, producing a pronounced strand-asymmetric substitution pattern (Figure 2C). Averaged across days 12 to 90, both transitions were roughly twice as frequent per nucleotide on the heavy strand in TOP3A KO compared to WT (C>T, 134.7 versus 66.7; A>G, 75.3 versus 31.7), whereas on the light strand the same classes were far rarer in both genotypes (C>T, 28.4 versus 13.1; A>G, 7.6 versus 3.1). The four transversion classes (A>C, A>T, C>A and C>G) occurred at substantially lower frequencies but showed a greater proportional increase in TOP3A KO iPSC-CMs, with a mean 2.9-fold elevation on the H strand, most marked for A>T (3.4-fold).

**Figure 2.**
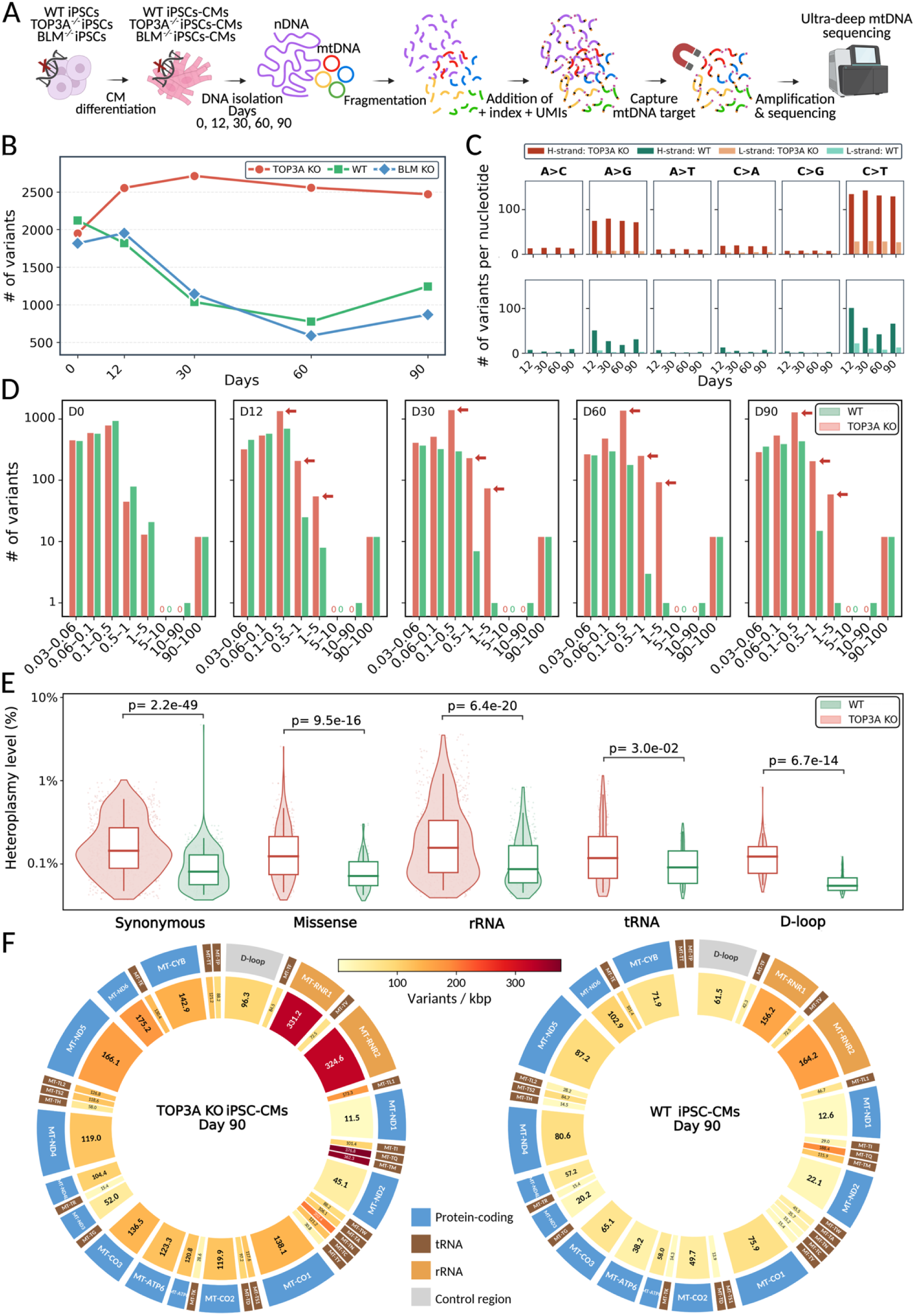
Ultra-deep mtDNA sequencing reveals increased mtDNA mutation load in TOP3A KO iPSC-CMs. **(A)** mtDNA-enriched ultra-deep sequencing workflow. Genomic DNA was isolated from WT, TOP3A KO, and BLM KO cells at day 0 and days 12, 30, 60, and 90 after cardiac differentiation. Fragmented DNA was ligated to adapters containing sample indices and UMIs, enriched by hybridization capture of mtDNA targets, amplified, and sequenced to approximately 500,000× mean raw mtDNA depth. **(B)** Number of distinct variants called per library across differentiation for TOP3A KO (red circles), WT (green squares), and BLM KO (blue diamonds). Each value represents the mutation load of one sequencing library. **(C)** Number of variants per corresponding reference nucleotide for the six substitution classes A>C, A>G, A>T, C>A, C>G, and C>T at days 12, 30, 60, and 90. Bars are separated by heavy (H)- and light (L)-strand assignment and genotype, with TOP3A KO in the upper row and WT in the lower row; dark and light shades denote H- and L-strand assignments, respectively. **(D)** Number of variants in TOP3A KO (red) and WT (green) within heteroplasmy intervals of 0.03%-0.06%, 0.06%-0.1%, 0.1%-0.5%, 0.5%-1%, 1%-5%, 5%-10%, 10%-50%, and 90%-100% at days 0, 12, 30, 60, and 90. The y-axis is logarithmic. Red arrows indicate intervals in which the number of TOP3A KO variants increased during differentiation. **(E)** Heteroplasmy levels of low-frequency variants (0.03%-10%) at day 90, grouped as synonymous, missense, rRNA, tRNA, or D-loop variants for TOP3A KO (red) and WT (green). Violins show the kernel density of log10-transformed heteroplasmy, clipped to the observed range and scaled according to the number of variants; boxes indicate the median and interquartile range, whiskers indicate the 5th and 95th percentiles, and points represent individual variant sites. Statistical comparisons were performed using two-sided Mann-Whitney U tests without adjustment for multiple comparisons. **(F)** Circos representation of variant density, expressed as variants per kilobase (variants/kbp), across mtDNA features at day 90 for TOP3A KO (left) and WT (right). The inner ring is shaded according to variant density, and the outer ring denotes protein-coding genes, tRNAs, rRNAs, and the control region.

Heteroplasmy refers to the fraction of mtDNA copies in a cell that carry a given variant^22^, measured here as allele frequency and reported as a percentage. Most variants occurred at low frequency, with levels between 0.03% and 0.5% (Figure 2D). During iPSC-CM maturation, the TOP3A KO distribution progressively shifted toward higher heteroplasmy categories. Variants in the 0.1%-0.5% interval increased from 787 at day 0 to 1,289 at day 90, with similar increases in the 0.5%-1% interval (44 to 205) and the 1%-5% interval (13 to 59). WT showed the opposite pattern, with variant counts declining across the corresponding intervals (935 to 432, 78 to 14, and 21 to 1). No variant in either genotype occupied the 5%-10% interval at any time point. Twelve variants in the 90%-100% heteroplasmy range were shared across all three isogenic lines at every time point, consistent with their common mitochondrial haplogroup H1c22. Coverage remained comparable across samples, supporting a genuine shift in heteroplasmy rather than altered detection sensitivity (Supplementary Figure 2). Specifically at day 90 post-differentiation, SNVs grouped by class of variation showed significantly higher heteroplasmy levels in TOP3A KO across coding (synonymous and missense) and non-coding mtDNA regions (rRNA, tRNA and D-loop) (Figure 2E). After normalization to gene length, the mitochondrial rRNA genes remained the most prominent mutational hotspots, with 331.2 SNVs/kbp in *MT-RNR1* and 324.6 SNVs/kbp in *MT-RNR2* in TOP3A KO, compared with 156.2 and 164.2 SNVs/kbp, respectively, in WT (Figure 2F). This was accompanied by a genome-wide accumulation of mtDNA SNVs in TOP3A KO, averaging 144.3 SNVs/kbp.

### Mutation load increase is driven by *de novo* variants arising mainly in mitochondrial ribosomal RNA genes

To determine the genetic basis of the increased mutation load in TOP3A KO iPSC-CMs, we applied rigorous filters (see Materials and Methods section), which resulted in the classification of SNVs as pre-existing (detected in all isogenic cell lines at every time point) or *de novo* (defined by absence from the iPSC baseline and from the isogenic control lines at all time points together with sustained detection at every time point following first appearance). We found that *de novo* SNVs accounted for the increase in TOP3A KO mutation load after differentiation (Figure 3A). Pre-existing SNVs remained detectable across the days post-differentiation in TOP3A KO, whereas in WT and BLM KO a defined subset of pre-existing SNVs were no longer detectable at certain time points. A fraction of these re-emerged at day 90, indicating that they had drifted to levels transiently indistinguishable from background.

**Figure 3.**
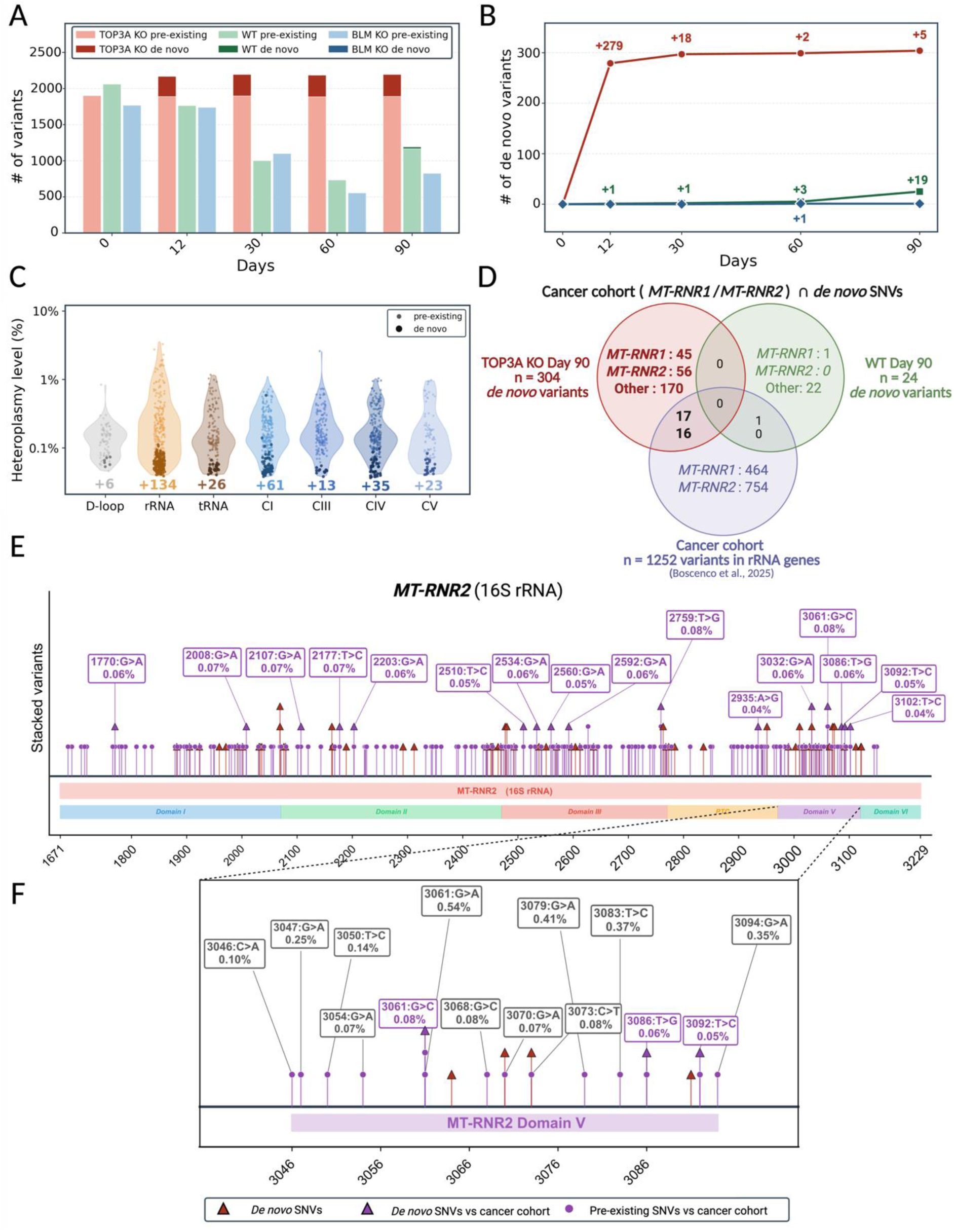
TOP3A-associated mtDNA variant accumulation is mainly driven by *de novo* SNVs enriched in mitochondrial rRNA genes. **(A)** Number of SNVs per library across days 0-90, partitioned into pre-existing (light shading) and *de novo* (dark shading) SNVs. TOP3A KO, WT, and BLM KO are shown in red, green, and blue, respectively. **(B)** Cumulative number of *de novo* SNVs across differentiation, using the same genotype color scheme. Numbers above the trajectories indicate newly acquired SNVs within each time interval. **(C)** Heteroplasmy distributions of pre-existing and *de novo* SNVs grouped by mitochondrial genomic category: D-loop, rRNA, tRNA, and OXPHOS complexes I, III, IV, and V. Violins show the heteroplasmy distributions, with individual SNV sites overlaid. Numbers below the violins indicate the number of *de novo* SNVs represented in each category. **(D)** Overlap between *de novo* SNVs in *MT-RNR1* and *MT-RNR2* detected at day 90 in TOP3A KO and WT cells and SNVs reported in *MT-RNR1* and *MT-RNR2* in the Genomics England pan-cancer mtDNA cohort (Boscenco et al., 2025). n indicates the total number of SNVs in each set. **(E)** Stacked-variant plot of *MT-RNR2* (16S rRNA; m.1671-3229), with its six secondary-structure domains and the peptidyl-transferase-associated region annotated below the gene. Red triangles denote *de novo* SNVs, purple triangles denote *de novo* SNVs also reported in the cancer cohort, and purple circles denote pre-existing SNVs also reported in the cancer cohort. Selected SNVs are labeled by nucleotide substitution and heteroplasmy level. **(F)** Magnified view of *MT-RNR2* Domain V (approximately m.3046-3094), using the same marker scheme. Selected SNVs are labeled by position, substitution, and heteroplasmy level.

The accumulation of *de novo* SNVs was most pronounced during the initial iPSC-to-iPSC-CM transition between days 0 and 12, with a total of 279 *de novo* SNVs acquired. This was followed by the appearance of additional *de novo* SNVs at later time points (18, 2, and 5 SNVs at days 30, 60, and 90, respectively), resulting in 304 *de novo* SNVs in total (Figure 3B). Thus, more than 90% of the *de novo* SNV load emerged during the first 12 days of differentiation, following the same transition-rich signature, enriched in the rRNA genes (Supplementary Figure 4A, B). Conversely, WT accumulated only 24 *de novo* SNVs over the entire time course, 19 of which were first detected at day 90, whereas BLM KO acquired only a single *de novo* SNV. To assess whether the TOP3A-associated mtDNA phenotype was reproducible across independently generated clones, we characterized two additional TOP3A KO clones (TOP3A KO clone 2 and clone 3) (Supplementary Figure 5A-F). TOP3A KO clone 2 was selected for further downstream analysis and subjected to ultra-deep mtDNA sequencing at days 0, 12, and 30 post-differentiation. TOP3A KO clone 2 showed an increase in total mtDNA mutation load together with early accumulation of *de novo* SNVs during differentiation (Supplementary Figure 5G and H), supporting a reproducible effect of TOP3A deficiency on mtDNA mutagenesis.

*De novo* SNVs were distributed throughout the mitochondrial genome and remained at low heteroplasmy levels across functional categories. The mitochondrial rRNA genes nevertheless represented the most prominent individual category, containing 134 *de novo* SNVs (Figure 3C; Supplementary Figure 4B). Protein-coding genes also contributed substantially, with 132 *de novo* SNVs distributed across mtDNA-encoded OXPHOS subunits: 61 in complex I, 13 in complex III, 35 in complex IV, and 23 in complex V (Supplementary Figure 6). Thus, although *de novo* SNV accumulation was genome-wide, *MT-RNR1* and *MT-RNR2* emerged as particularly prominent sites, consistent with recent evidence identifying mitochondrial rRNA genes as recurrent sites of somatic mtDNA variation in cancer^23^.

Comparison with the Genomics England pan-cancer mtDNA cohort further revealed substantial overlap with SNVs observed across tumors. Among the TOP3A KO *de novo* SNVs, 33 mitochondrial rRNA SNVs were also identified in the cancer cohort as exact position- and variation-matched events, comprising 17 in *MT-RNR1* and 16 in *MT-RNR2* (Figure 3D; Supplementary Figure 7). Of these, only *MT-RNR1* m.933G>A was additionally classified by Boscenco et al. as a statistically significant recurrent SNV hotspot. Extending the analysis to the full day 90 rRNA SNV landscape (including pre-existing variants), substantially increased the overlap with the cancer cohort (Supplementary Figure 8A). A total of 185 SNVs in *MT-RNR2* matched cohort variants, including 169 pre-existing and 16 *de novo* SNVs (Figure 3E). These overlapping variants were distributed across all six secondary-structure domains, with enrichment in domain V, where several reached the highest heteroplasmy levels (Figure 3F). An additional 99 *MT-RNR1* SNVs overlapped the cohort, comprising 82 pre-existing and 17 *de novo* SNVs (Supplementary Figure 8C).

We next restricted the analysis to the 138 positions identified by Boscenco et al. as statistically significant recurrent SNV hotspots. Four TOP3A KO *de novo* SNVs mapped to these positions: *MT-RNR1* m.933G>A and m.1336G>T, *MT-RNR2* m.3091G>T, and *MT-ND4* m.10914G>A (Supplementary Figure 8B). At m.933 and m.10914, both position and nucleotide substitution matched the recurrent cancer hotspot, whereas the TOP3A KO SNVs at m.1336 and m.3091 affected the same hotspot positions but differed from the recurrent G>A substitutions reported in the cancer cohort. Structural analysis by Boscenco et al. localized m.933 and m.1336 to a cluster of recurrent *MT-RNR1* hotspots at the entrance to the mitochondrial small ribosomal subunit (mtSSU) mRNA channel. More broadly, recurrent mitochondrial rRNA hotspots mapped to defined mitoribosomal structural features and to loci under strong germline purifying selection, supporting potential loss-of-function consequences at these positions^23^.

Together, these findings identify the mitochondrial rRNA genes as major sites of *de novo* SNV accumulation following TOP3A loss and reveal two levels of convergence with cancer-associated mtDNA variation: broad overlap with SNVs observed across tumors and a more selective convergence on statistically significant recurrent hotspots, including structurally relevant mitoribosomal loci.

### TOP3A deficiency impairs mtDNA copy number expansion and reshapes mitochondrial DNA variant heteroplasmy dynamics

Given the marked accumulation of *de novo* SNVs in TOP3A KO during early differentiation, we next assessed whether differences in mtDNA copy number could contribute to the observed variant burden and heteroplasmy patterns. Using PacBio HiFi long-read whole-genome sequencing (LR WGS), we estimated mtDNA copy number at days 12 and 60 directly from mitochondrial and nuclear sequencing depth within the same libraries. We estimated mtDNA copy number relative to four independent diploid nuclear reference loci (*ALB*, *B2M*, *RPP30*, and *TBP*). At day 12, WT and TOP3A KO showed comparable mtDNA copy numbers of approximately 50 copies per cell. By day 60, however, WT iPSC-CMs reached approximately 1,084 copies per cell, whereas TOP3A KO iPSC-CMs reached only approximately 96 copies per cell, corresponding to an 11.3-fold difference and demonstrating impaired mtDNA copy-number expansion during iPSC-CM maturation (Figure 4A).

**Figure 4.**
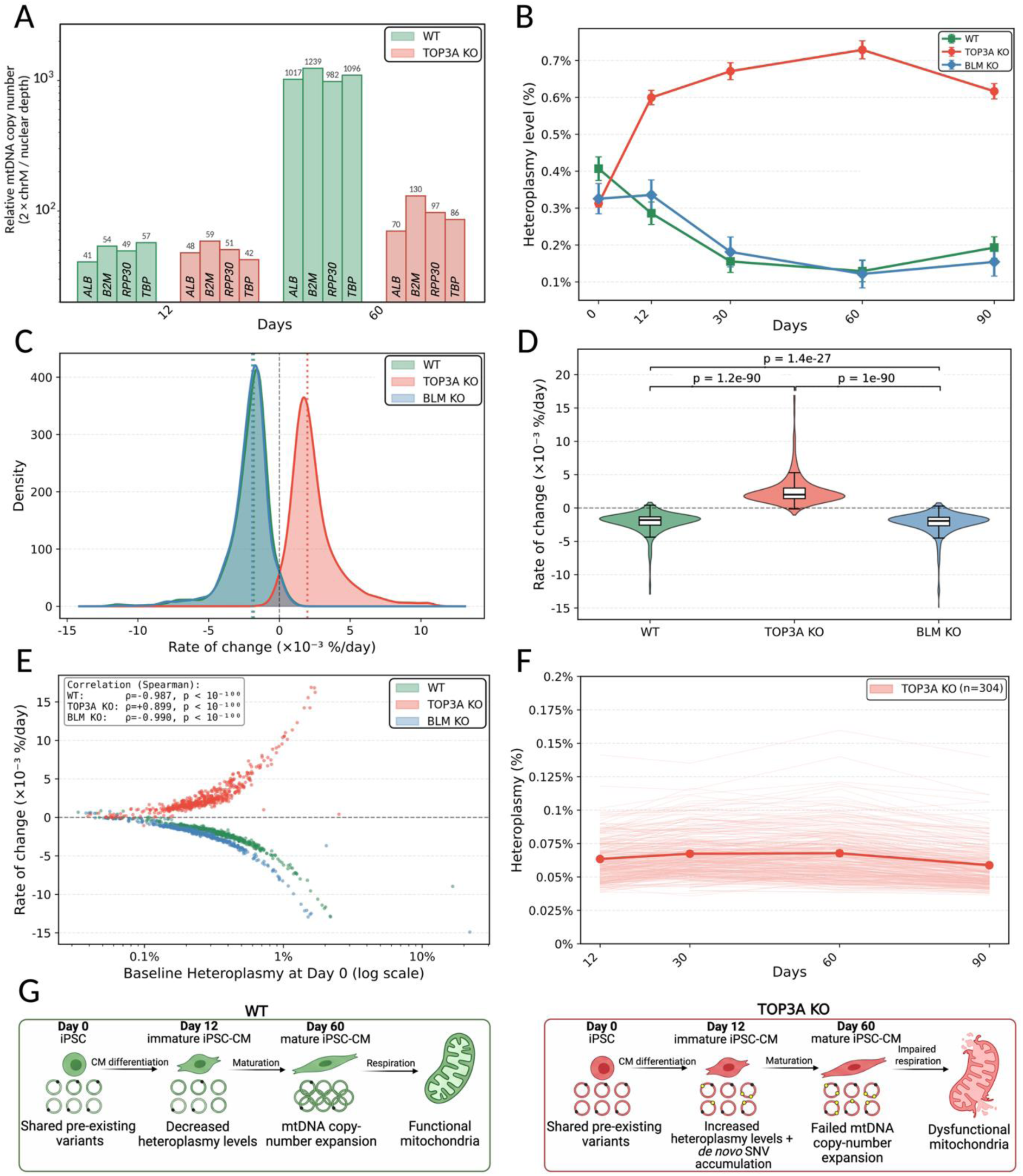
TOP3A deficiency impairs copy-number expansion during iPSC-CM maturation and reshapes pre-existing mtDNA heteroplasmy dynamics. **(A)** Relative mtDNA copy number estimated from PacBio HiFi whole-genome sequencing generated at approximately 30× nuclear depth. Estimates were calculated as twice the mean chrM depth divided by the mean depth of each diploid nuclear reference locus (*ALB*, *B2M*, *RPP30*, and *TBP*). Bars represent estimates derived independently from each nuclear locus. One library was analyzed per genotype and time point. WT and TOP3A KO are shown in green and red, respectively. **(B)** Mean heteroplasmy across days 0-90 for 545 variant positions detected at every time point in all three genotypes after exclusion of variants with a maximum allele frequency of ≥90%. WT, TOP3A KO, and BLM KO are shown in green, red, and blue, respectively. **(C)** Kernel-density distributions of the per-variant heteroplasmy rate of change for WT (green), TOP3A KO (red), and BLM KO (blue), calculated as the slope of an ordinary least-squares regression of heteroplasmy against time and expressed as ×10^-^³ %/day. Dotted vertical lines indicate genotype medians, and the dashed vertical line indicates zero change. **(D)** Rate of heteroplasmy change for the 545 matched variant positions, shown as violins with overlaid boxplots. Matched-position comparisons were performed using Wilcoxon signed-rank tests. **(E)** Relationship between heteroplasmy rate of change and baseline heteroplasmy at day 0 for WT (green), TOP3A KO (red), and BLM KO (blue), displayed on a logarithmic x-axis. Each point represents one variant position, and associations were assessed using Spearman rank correlation. **(F)** Heteroplasmy trajectories of the 304 TOP3A KO *de novo* SNVs from days 12-90. Thin lines represent individual SNVs, and the bold line and shading represent mean ± SEM. **(G)** Schematic summary of altered mtDNA maintenance following TOP3A loss during iPSC-CM differentiation and maturation. WT and TOP3A KO are represented in green and red, respectively. WT and TOP3A KO cells carry shared pre-existing heteroplasmic mtDNA variants at the iPSC stage (day 0, D0). In WT cells, heteroplasmy levels decrease during differentiation (day 12, D12), followed by mtDNA copy-number expansion and mitochondrial maturation (day 60, D60). In TOP3A KO cells, differentiation is accompanied by accumulation of low frequency *de novo* mtDNA SNVs and progressive increases in the heteroplasmy of shared pre-existing variants. This altered mtDNA landscape is followed by impaired mtDNA copy-number expansion and mitochondrial dysfunction during iPSC-CM maturation. Black circles indicate pre-existing heteroplasmic variants, and red circles indicate low frequency *de novo* SNVs.

To independently validate these findings and resolve mtDNA copy-number dynamics during early differentiation, we quantified relative mtDNA/nuclear DNA (nDNA) copy number by qPCR at days 0, 2, 6, 10, 12, and 60. Relative copy number remained comparable between WT and TOP3A KO throughout early differentiation, including at day 12, whereas a genotype-specific divergence emerged by day 60, when WT iPSC-CMs increased mtDNA copy number approximately 2.4-fold while TOP3A KO failed to undergo a comparable expansion (Supplementary Figure 9A). Importantly, similar relative copy number through day 12 indicates that the early accumulation of 279 *de novo* SNVs was not driven by differences in mtDNA pool size.

Given the earlier shift toward higher heteroplasmy categories in TOP3A KO (Figure 2D), we examined the trajectories of pre-existing variants in greater detail across the isogenic cell lines, restricting the analysis to 545 variant positions detected at every time point in all three genotypes. Mean heteroplasmy increased progressively in TOP3A KO from 0.31% at day 0 to 0.73% by day 60, whereas it decreased in WT from 0.41% to 0.13% and in BLM KO from 0.33% to 0.12% over the same interval (Figure 4B). A partial recovery was observed by day 90, reaching 0.19% in WT and 0.15% in BLM KO. These trajectories are consistent with the reduction in total variant calls observed in the control lines, as variants declining below the detection threshold would no longer be retained as confident calls.

Consistent with these opposing trajectories, heteroplasmy rates of change were predominantly positive in TOP3A KO (median +2.01 × 10^-^³ %/day) and negative in WT and BLM KO (medians -1.81 and -1.94 × 10^-^³ %/day, respectively; Figure 4C and D). We next examined whether pathogenicity, nucleotide substitution class, genomic category, or baseline iPSC heteroplasmy was associated with the rate of heteroplasmy change. No strong association was observed with predicted pathogenicity (r = -0.23; Supplementary Figure 9B) or nucleotide substitution class (Supplementary Figure 9C), although rRNA variants showed the highest average rates of change (Supplementary Figure 9D). Baseline heteroplasmy showed the strongest association, but in opposite directions between genotypes: variants present at higher heteroplasmy at day 0 increased most rapidly in TOP3A KO (Spearman ρ = +0.899) and decreased rapidly in WT and BLM KO (ρ = -0.987, ρ = -0.990; Figure 4E). Stratification by baseline heteroplasmy and correlation analyses further supported baseline allele frequency as the principal correlate of heteroplasmy trajectory (Supplementary Figure 9E and F).

Thus, TOP3A deficiency alters the frequency-dependent behavior of pre-existing mtDNA variants during cardiomyocyte differentiation and maturation. In contrast to these pre-existing variants, the mean heteroplasmy of the 304 TOP3A KO *de novo* SNVs remained stable at approximately 0.06%-0.07% from day 12 to day 90 (Figure 4F), indicating persistent low-frequency variation without detectable clonal expansion over the subsequent maturation period. Together, these findings support a model in which TOP3A loss produces an early burst of low-frequency *de novo* mtDNA mutagenesis independently of mtDNA copy-number changes, followed by altered heteroplasmy trajectories of pre-existing variants and a later failure of mtDNA copy-number expansion during cardiomyocyte maturation (Figure 4G).

## Discussion

Biallelic loss-of-function variants in *TOP3A* cause a clinical spectrum ranging from BSyn to clinical mitochondriopathy that can also include cardiomyopathy^5^, yet the pathogenesis of *TOP3A*-associated cardiomyopathy is unknown. Using an isogenic iPSC-CM model, we demonstrated that TOP3A deficiency impairs OXPHOS function. At the molecular level, this was accompanied by a differentiation-triggered accumulation of low-frequency *de novo* mtDNA SNVs, a progressive increase in heteroplasmy levels of variants shared across isogenic lines and an impaired mtDNA copy-number expansion during iPSC-CM maturation. To our knowledge, this study provides the first human iPSC-CM model of TOP3A deficiency and directly connects defective mtDNA maintenance with mitochondrial dysfunction. More broadly, it establishes mtDNA instability as a plausible initiator and contributor to *TOP3A*-associated cardiomyopathy and supports investigation of mtDNA mutational signatures in other genetic forms of inherited cardiomyopathy.

### Mitochondrial dysfunction is caused by the combined consequences of TOP3A-related mtDNA instability

We demonstrated that loss of TOP3A leads to mitochondrial dysfunction in mature day 60 iPSC-CMs. These findings are supported by the established role of TOP3A within mitochondria in resolving the hemicatenane formed at the origin of H-strand replication during mtDNA replication termination and in allowing proper nucleoid segregation^3^. TOP3A additionally supports replication-fork progression and sustains steady-state mitochondrial transcript levels^8^. Previous experimental models have likewise linked TOP3A loss to mitochondrial dysfunction, including CRISPR-engineered TOP3A KO HCT116 cells and a Drosophila model lacking the mitochondrial top3α isoform, which exhibited decreased mitochondrial membrane potential and ATP content^9,24^. More recently, homozygous *Top3a* mutant embryonic stem cells were shown to develop progressive mtDNA depletion accompanied by increased ROS levels^25^. Nevertheless, these studies primarily established the functional consequences of TOP3A loss without defining the mtDNA sequence-level changes that connect impaired genome stability to mitochondrial dysfunction. In the present study, we show that *de novo* variants and most pre-existing variants can be detected at low frequencies, which lie far below the classical heteroplasmic thresholds for biochemical expression, and no single variant is likely to account for the mitochondrial OCR defect we observed. The functional phenotype is instead best understood as the combined consequence of the additive functional burden of accumulating low-frequency as well as persisting pre-existing variants, along with defective copy-number expansion and described larger mtDNA deletions^3,4,26–28^.

### Cardiac differentiation promotes *de novo* SNV accumulation in the absence of TOP3A

TOP3A KO accumulated most *de novo* variants between day 0, at the iPSC stage, and day 12, when cells had committed to the cardiac lineage. Cardiac commitment changes the energy use towards mitochondrial production and also increases energy demand and this drives extensive mitochondrial remodeling, including greater OXPHOS dependence, mitochondrial biogenesis, and changes in substrate use^29–32^. These processes increase mtDNA replication and transcription while remodeling mitochondrial nucleoids, creating a vulnerable window in which TOP3A loss promotes mtDNA instability particularly up to day 12. The proofreading-deficient polymerase gamma (Polg) mutator mouse provides a useful comparison with TOP3A deficiency. Homozygous *Polg* mutator mice accumulate mtDNA SNVs and deletions together with mitochondrial dysfunction and premature aging phenotypes^33–35^. C>T transitions also predominate in the Polg mutator mouse, where replication errors accumulate without Polg 3′-5′ exonuclease proofreading^36^. Both deficiencies therefore produce transition-rich low-frequency SNV spectra consistent with spontaneous base deamination during strand-asymmetric replication. However, TOP3A has no established role in proofreading nascent DNA, so the true mechanism of mutagenesis in TOP3A KO iPSC-CMs remains unknown. Additionally, concerns are frequently raised regarding the authenticity of what some studies, including the present study, define as *de novo* mtDNA mutations. At 500,000× coverage, manual BAM inspection revealed sparse reads at nearly every mtDNA position, making low-level day 0 support insufficient to establish a genuine baseline variant. Although LoFreq, UMI consensus error correction, and downstream filtering increase confidence in our calls, we cannot completely exclude the presence of these variants below the day 0 detection threshold, although this appears unlikely given the sequencing depth and stringent filtering strategy. Our SNV-focused pipeline also cannot assess large-scale mtDNA deletions, which represent another relevant component of TOP3A-associated instability^4,37^, but these have been characterized previously and were not the focus of the present study.

### Mitochondrial rRNA genes represent prominent hotspots of mtDNA instability in TOP3A KO cells

We identified the mitochondrial rRNA genes as prominent sites of mtDNA variant accumulation, encompassing both pre-existing variants and approximately 45% of all *de novo* variants. Pathogenic variants within mitochondrial rRNA genes have previously been associated with mitochondrial and cardiac disease, including a *de novo MT-RNR2* variant associated with combined respiratory-chain deficiency and myopathy^38^ and the m.2336T>C *MT-RNR2* variant linked to hypertrophic cardiomyopathy^39,40^. The distinctive location of the rRNA loci may contribute to variant enrichment in *MT-RNR1* and *MT-RNR2*, as these genes lie immediately downstream of the control region and near OriH, placing them where transcription initiation and replication termination intersect. TOP3A loss could therefore increase topological stress across the adjacent rRNA loci and promote local instability^41,42^, although we did not directly measure topological stress in these iPSC-CMs. As described above, the pan-cancer cohort revealed recurrent somatic hotspots in *MT-RNR1* and *MT-RNR2* at conserved positions involved in mitoribosome interactions^23^. The authors further demonstrated that these mt-rRNA variants were subject to positive selection and could act functionally dominantly by impairing mitochondrial translation. Notably, they demonstrated that the m.1227G>A *MT-RNR1* variant, which was also detected among our pre-existing variants, impaired mitochondrial function and reduced respiratory-chain subunit abundance from approximately 10% heteroplasmy, below conventional biochemical thresholds for mtDNA pathogenicity. Although individual variants in our model remained below this level, their cumulative burden may contribute to the mitochondrial dysfunction observed in TOP3A KO iPSC-CMs, a possibility that will require further experimental validation.

### TOP3A deficiency alters the selection pressures exerted upon pre-existing mtDNA heteroplasmies

At the iPSC level at day 0, 1,735 mtDNA SNV sites were shared across the WT, TOP3A KO and BLM KO lines, indicating that these variants were already present before the individual isogenic lines were established (Supplementary Figure 2C). These pre-existing variants may be maternally inherited heteroplasmies and somatic mosaic variants already present in the WT fibroblasts, which became enriched during iPSC reprogramming^43–45^. A subset may also have arisen *de novo* during reprogramming or during expansion of the parental WT iPSC line before genome editing^46^, consistent with the substantial overlap of these pre-existing variants with somatic mtDNA SNVs independently acquired in the pan-cancer cohort (Figure 3E; Supplementary Figure 8C). Subsequent clonal isolation and culture may have further altered heteroplasmy levels, as substantial differences in mtDNA heteroplasmy have been reported between independently derived iPSC clones originating from the same parental cell population^44,47^.

The pre-existing variants displayed divergent heteroplasmy trajectories, with mean heteroplasmy increasing in TOP3A KO cells but declining in WT and BLM KO cells. The decline in the control lines could reflect purifying selection during differentiation, consistent with the progressive depletion of pathogenic mtDNA variants reported in differentiating hematopoietic and immune lineages^48–50^. Our findings raise the possibility that selection pressures during the iPSC-to-iPSC-CM transition, previously demonstrated for functionally consequential high-heteroplasmy mtDNA variants, may also extend to low-frequency variants of uncertain functional significance. By contrast, TOP3A deficiency may have altered this selective environment and narrowed the pool of cells capable of completing cardiac differentiation, allowing subpopulations carrying higher levels of pre-existing variants to contribute disproportionately, a mechanism that has been reported before^51^. Further single-cell analyses might help to determine whether the observed heteroplasmy shifts occur consistently across individual cardiomyocytes or are concentrated within a subset of cells, and whether they involve the broader intracellular mtDNA pool within a cell, or only a fraction of mtDNA molecules.

Our findings therefore suggest that mtDNA maintenance is an essential determinant of cardiac health. Analysis of mtDNA sequence integrity, associated heteroplasmy levels and cell-specific copy number of mtDNA molecules should therefore form a core component of inherited cardiomyopathy research and diagnostics, with the potential to further clarify the molecular pathogenesis of otherwise unexplained diseases. Despite its importance, mtDNA maintenance and integrity are still being explored, highlighting the need for ultra-deep mtDNA sequencing strategies to reveal both disease-associated alterations and fundamental genetic processes governing the emergence, persistence and remodeling of mtDNA variation. In addition to inherited cardiomyopathy, our experimental setup could enable the investigation of mechanisms contributing to the accumulation of age-associated somatic mtDNA mutations in various organ systems including the heart, although this application will require validation across additional sequencing libraries and extended culture durations.

## Acknowledgments

We gratefully acknowledge Karin Boss for critical proofreading of the manuscript, and Christian Müller for excellent technical assistance. This work was supported by the German Research Foundation (DFG, Deutsche Forschungsgemeinschaft) under Germany’s Excellence Strategy (EXC 2067/1-390729940) to BW. BW acknowledges further funding by DZKJ (German Center for Child and Adolescent Health) and DZHK (German Centre for Cardiovascular Research, partner site Lower Saxony). This work was partly accomplished within the Center for Undiagnosed Congenital Syndromes and Clinical Genome Medicine at the Center of Rare Diseases Göttingen (ZSEG). MG was supported by the Ph.D. program ‘Genome Science’ - International Max Planck Research School.

## Disclosures

None.

## Supplemental Material

Figures S1-S9

## Notes

### Competing Interest Statement

The authors have declared no competing interest.

