## Supplemental_figures for "Distinct mitochondrial DNA single-nucleotide variant signatures in *TOP3A*-deficient cardiomyocytes"

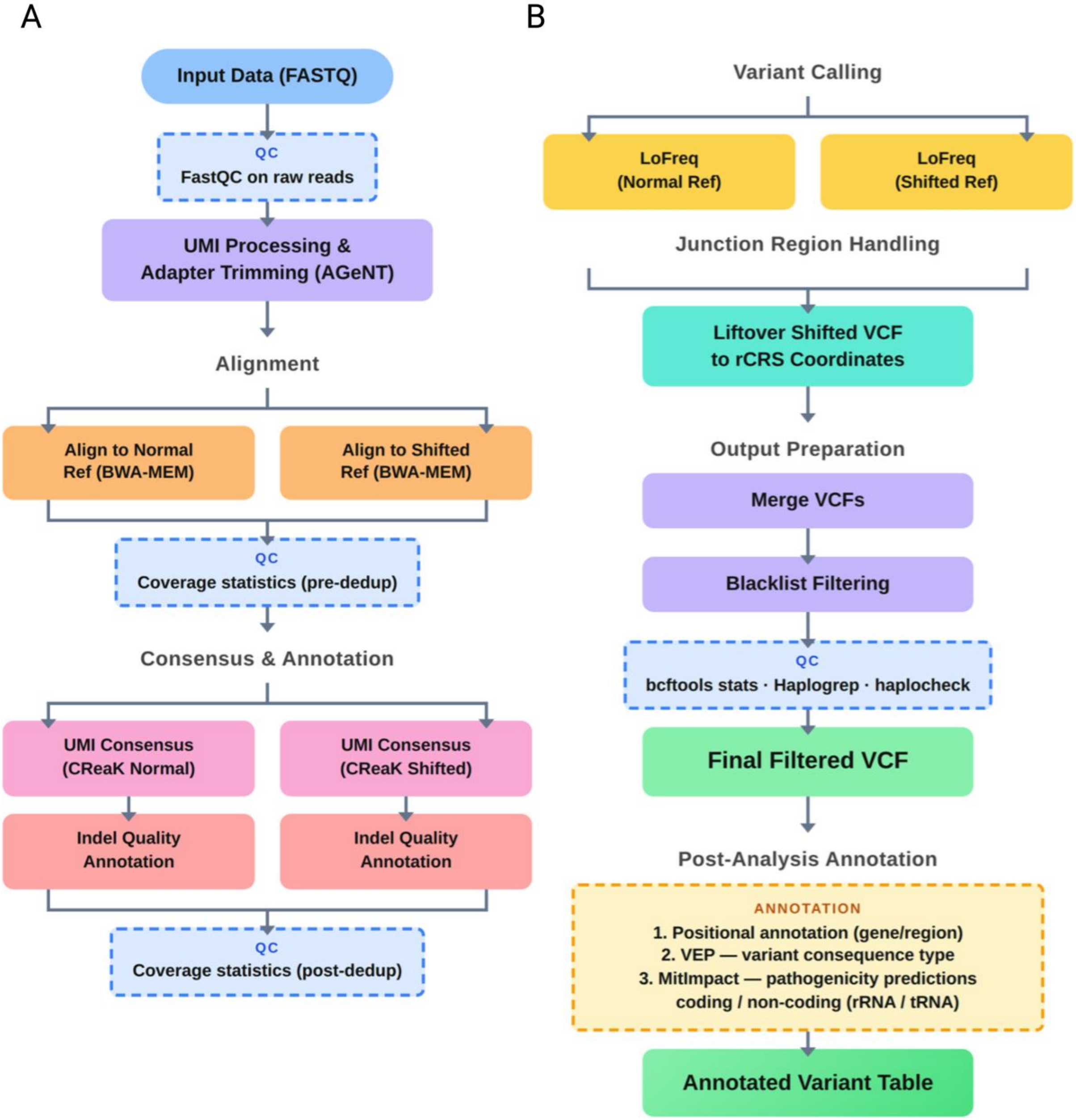


**Supplementary Figure 1. mtDNA variant-calling pipeline. (A)** Preprocessing, alignment, and consensus generation. Raw FASTQ files underwent quality control using FastQC, UMI extraction and adapter trimming using AGeNT, and parallel alignment with BWA-MEM to the standard and shifted revised Cambridge Reference Sequence (rCRS). Coverage was assessed before deduplication, UMI-consensus reads were generated for each alignment using CReaK, indel quality was annotated, and post-deduplication coverage was assessed. **(B)** Variant calling, merging, and annotation. Variants were called independently from the standard and shifted alignments using LoFreq. Calls from the shifted reference were lifted to rCRS coordinates, merged, filtered against blacklisted positions, and evaluated using bcftools stats, Haplogrep, and haplocheck. The final variant set was annotated for genomic position, gene or region, predicted consequence using Ensembl Variant Effect Predictor, and pathogenicity using MitImpact.

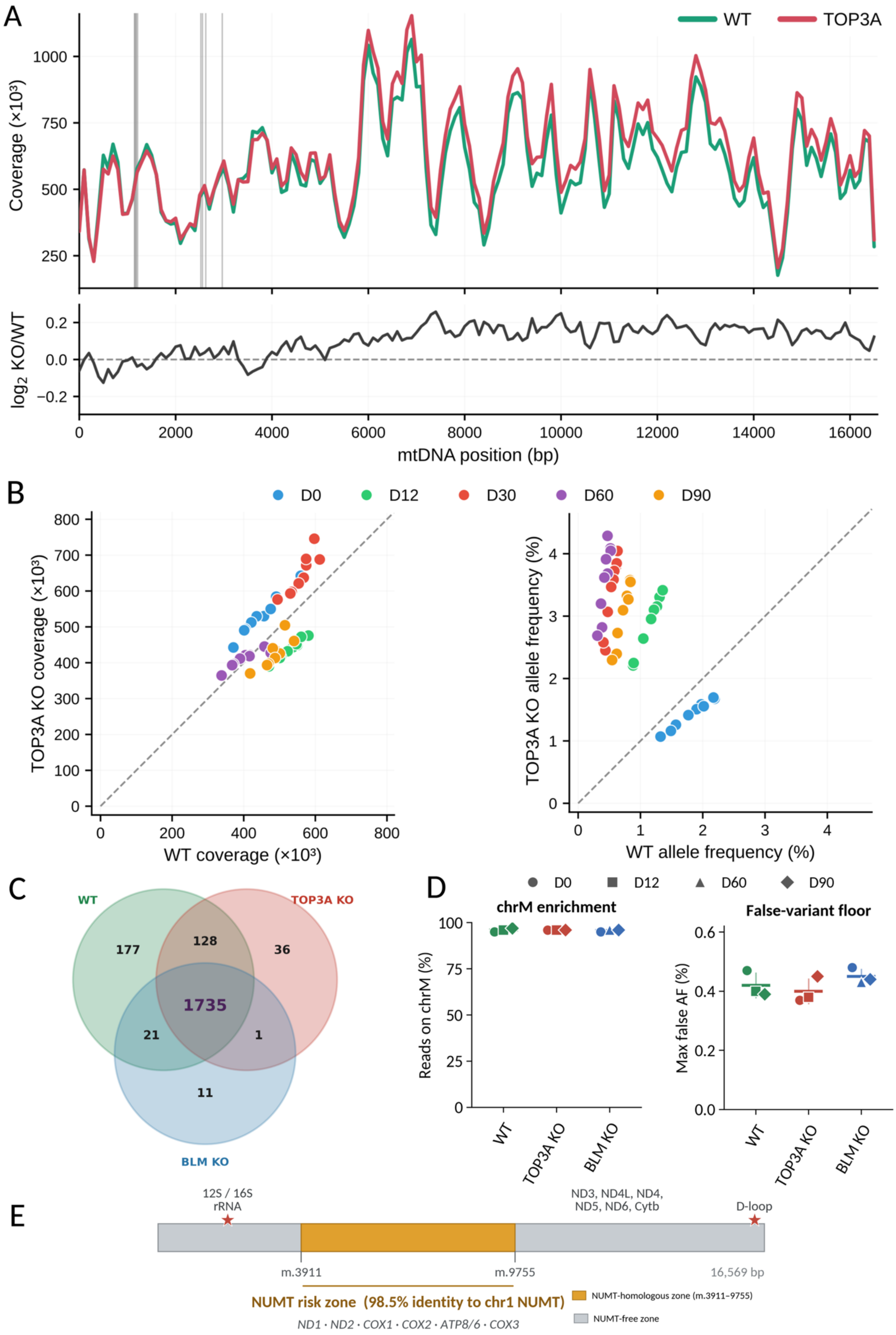

A
WT
TOP3A
Coverage (×10³)
log₂ KO/WT
mtDNA position (bp)
B
D0
D12
D30
D60
D90
TOP3A KO coverage (×10³)
WT coverage (×10³)
TOP3A KO allele frequency (%)
WT allele frequency (%)
C
WT
TOP3A KO
BLM KO
177
128
36
1735
21
1
11
D
chrM enrichment
Reads on chrM (%)
False-variant floor
Max false AF (%)
WT
TOP3A KO
BLM KO
E
12S / 16S
rRNA
ND3, ND4L, ND4,
ND5, ND6, Cytb
D-loop
m.3911
m.9755
16,569 bp
NUMT risk zone (98.5% identity to chr1 NUMT)
ND1 · ND2 · COX1 · COX2 · ATP8/6 · COX3
NUMT-homologous zone (m.3911–9755)
NUMT-free zone

**Supplementary Figure 2. Sequencing-depth comparability and assessment of NUMT-derived artifacts. (A)** Per-position UMI-consensus mtDNA coverage, expressed in thousands of reads, across the mitochondrial genome for WT (green) and TOP3A KO (red). Lines denote the mean across the five differentiation time points (days 0, 12, 30, 60, and 90), and gray vertical bands indicate blacklisted or excluded positions. The lower panel shows the $\log_2$ TOP3A KO/WT coverage ratio across the mitochondrial genome; the dashed horizontal line indicates a ratio of 1 ($\log_2$ ratio = 0). **(B)** Comparison of WT and TOP3A KO UMI-consensus BAM depth (left) and BAM-level allele frequency (right) at nine representative variant positions across five time points (45 paired observations). Points are colored by time point, and dashed lines indicate identity. The mean TOP3A KO/WT coverage ratio was $1.015 \pm 0.151$, whereas the mean TOP3A KO/WT allele-frequency ratio was $4.42 \pm 2.70$. Ratios were compared with a null value of 1 using one-sample t-tests and Wilcoxon signed-rank tests. **(C)** Three-way Venn diagram of mtDNA SNV calls at day 0 for WT (green), TOP3A KO (red), and BLM KO (blue); indels were excluded. A total of 1,735 SNVs were shared across all three isogenic lines. **(D)** Sequencing quality-control metrics for WT (green), TOP3A KO (red), and BLM KO (blue). Point shape denotes sampling day (days 0, 12, 60, and 90). Metrics comprise the percentage of reads mapping to chromosome M (chrM enrichment) and the maximum estimated false allele frequency. **(E)** Linear representation of the 16,569-bp mitochondrial genome. The orange interval denotes the NUMT-homologous region at m.3911–9755, which shares 98.5% sequence identity with the chromosome 1 NUMT and spans *MT-ND1* through *MT-CO3*; flanking regions are NUMT-free. Red stars indicate the 12S/16S rRNA and D-loop regions containing elevated TOP3A KO variant levels outside the NUMT-homologous interval.

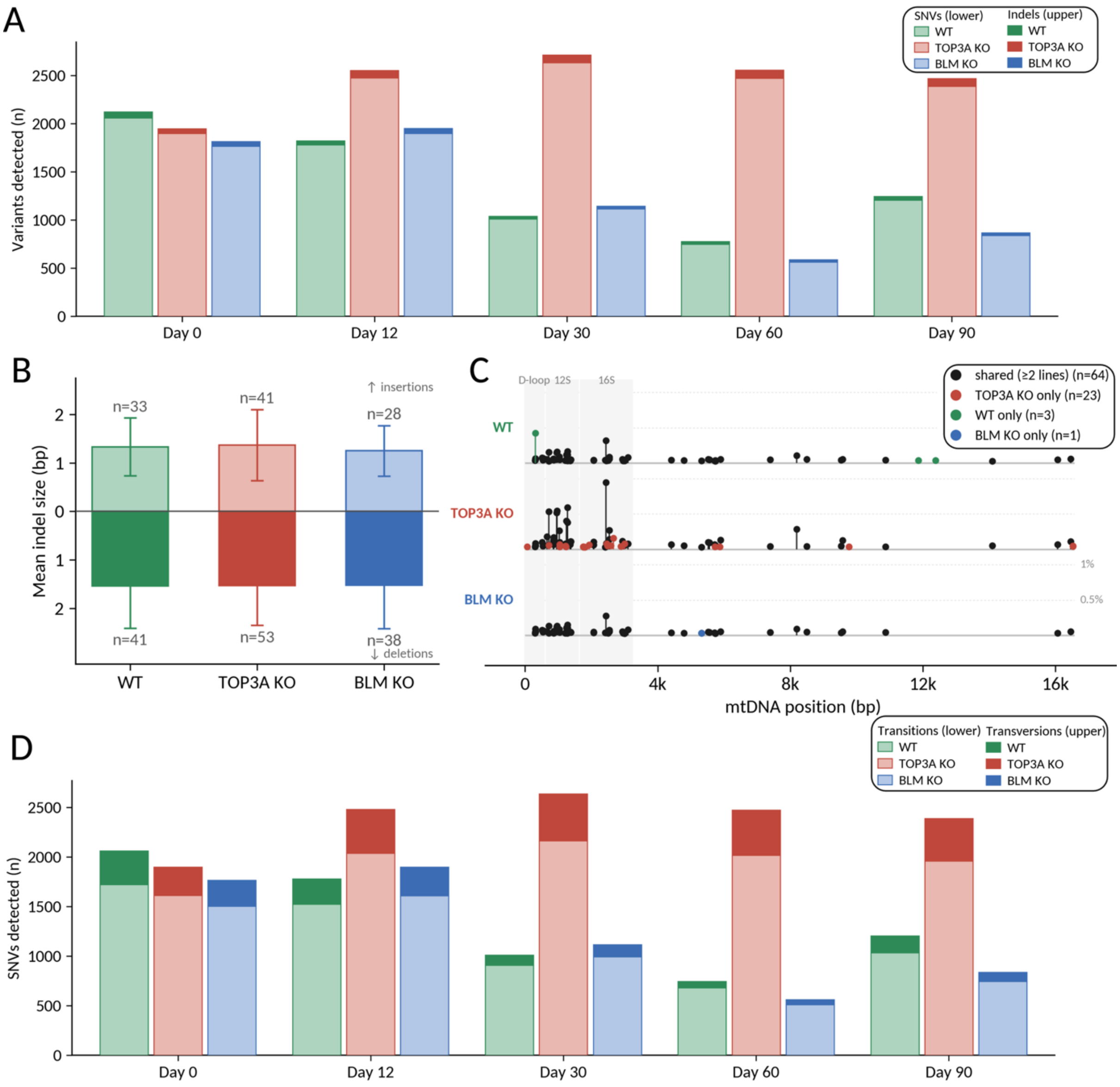


**Supplementary Figure 3. mtDNA SNV, indel, and substitution spectra across WT, TOP3A KO, and BLM KO.** **(A)** Total mutation load at days 0, 12, 30, 60, and 90, shown as stacked bars with SNVs in the lower, lighter portion and indels in the upper, darker portion. Within each time point, WT, TOP3A KO, and BLM KO are shown from left to right in green, red, and blue, respectively. **(B)** Mean indel size, with insertions plotted upward and deletions downward. Bars indicate the mean, whiskers indicate ± SD, and n denotes the number of distinct indel variants in each direction. **(C)** Genome-wide indel landscape. Each distinct indel is represented as a needle at its mtDNA coordinate on the corresponding genotype track, with needle height indicating mean heteroplasmy. Black denotes variants shared by ≥2 genotypes, whereas red, green, and blue denote variants private to TOP3A KO, WT, and BLM KO, respectively. The D-loop, *MT-RNR1* (12S rRNA), and *MT-RNR2* (16S rRNA) intervals are shaded; the homoplasmic m.3105/16S reference-alignment artifact was excluded. **(D)** Substitution spectrum over the same time course, with transitions (A↔G and C↔T) shown in the lower portion and transversions in the upper portion. Genotype ordering and colors are as in **(A)**.

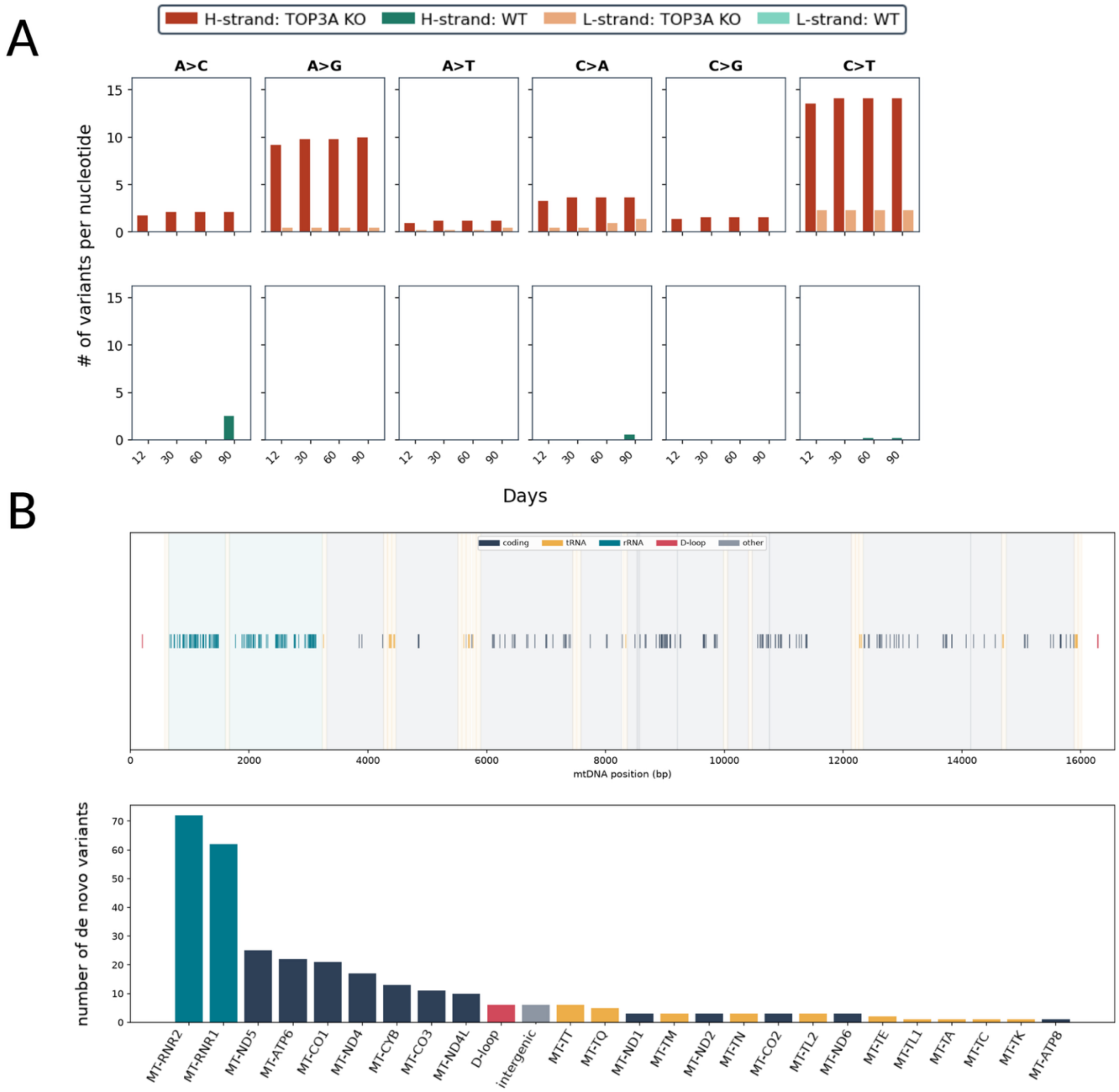


**Supplementary Figure 4. TOP3A KO *de novo* mtDNA SNVs show a transition-rich mutational signature and are enriched in *MT-RNR1* and *MT-RNR2*. (A)** Number of SNVs per corresponding reference nucleotide for the six substitution classes A>C, A>G, A>T, C>A, C>G, and C>T at days 12, 30, 60, and 90. Bars are separated by heavy (H)- and light (L)-strand assignment and genotype, with TOP3A KO shown in the upper row and WT in the lower row. Dark red and light red denote TOP3A KO H- and L-strand assignments, respectively, whereas dark green and light green denote the corresponding WT assignments. **(B)** Genomic distribution of the 304 *de novo* SNVs identified in TOP3A KO iPSC-CMs. Upper, position of each *de novo* SNV along the mitochondrial genome, colored according to genomic annotation. Lower, number of *de novo* SNVs per gene or genomic region, ordered by decreasing SNV count. Protein-coding regions are shown in dark blue, tRNAs in orange, rRNAs in teal, the D-loop in red, and intergenic or other regions in gray. Each SNV site was counted once according to its genomic position.

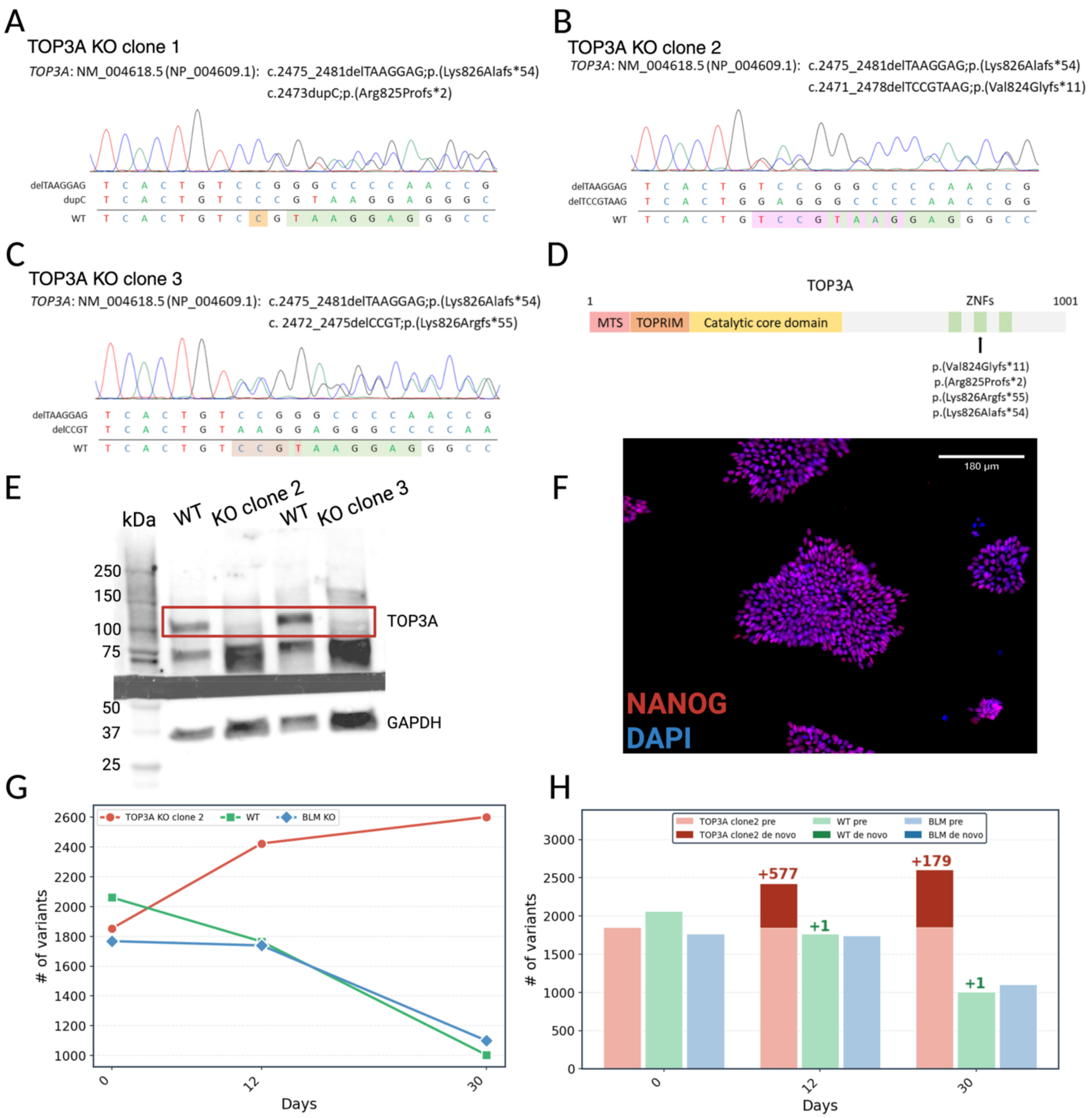


**Supplementary Figure 5. An independent TOP3A KO clone reproduces early mtDNA *de novo* variant accumulation. (A–C)** Sanger sequencing chromatograms of TOP3A KO clones 1 **(A)**, 2 **(B)**, and 3 **(C)** at the edited *TOP3A* locus, annotated relative to transcript NM_004618.5 and protein NP_004609.1. The edited alleles and WT reference sequence are aligned below each trace. Clone 1 represents the primary TOP3A KO clone used throughout the study. **(D)** Schematic representation of the TOP3A protein showing the positions of the frameshift variants identified across the three TOP3A KO clones. MTS, mitochondrial targeting sequence; TOPRIM, topoisomerase–primase domain; ZNFs, zinc-finger domains. **(E)** Immunoblot analysis of TOP3A in WT and TOP3A KO clones 2 and 3. GAPDH was used as a loading control. **(F)** Representative immunofluorescence image of TOP3A KO clone 2 iPSCs stained for the pluripotency marker NANOG (red) and DAPI (blue). Scale bar, 180 µm. **(G)** Number of mtDNA variants detected at days 0, 12, and 30 in TOP3A KO clone 2 (red), WT (green), and BLM KO (blue), demonstrating reproduction of the early increase in mtDNA variant burden observed with the primary TOP3A KO clone. **(H)** Corresponding mtDNA variant counts partitioned into pre-existing (light shading) and *de novo* (dark shading) variants, showing the accumulation of *de novo* variants in TOP3A KO clone 2 during differentiation.

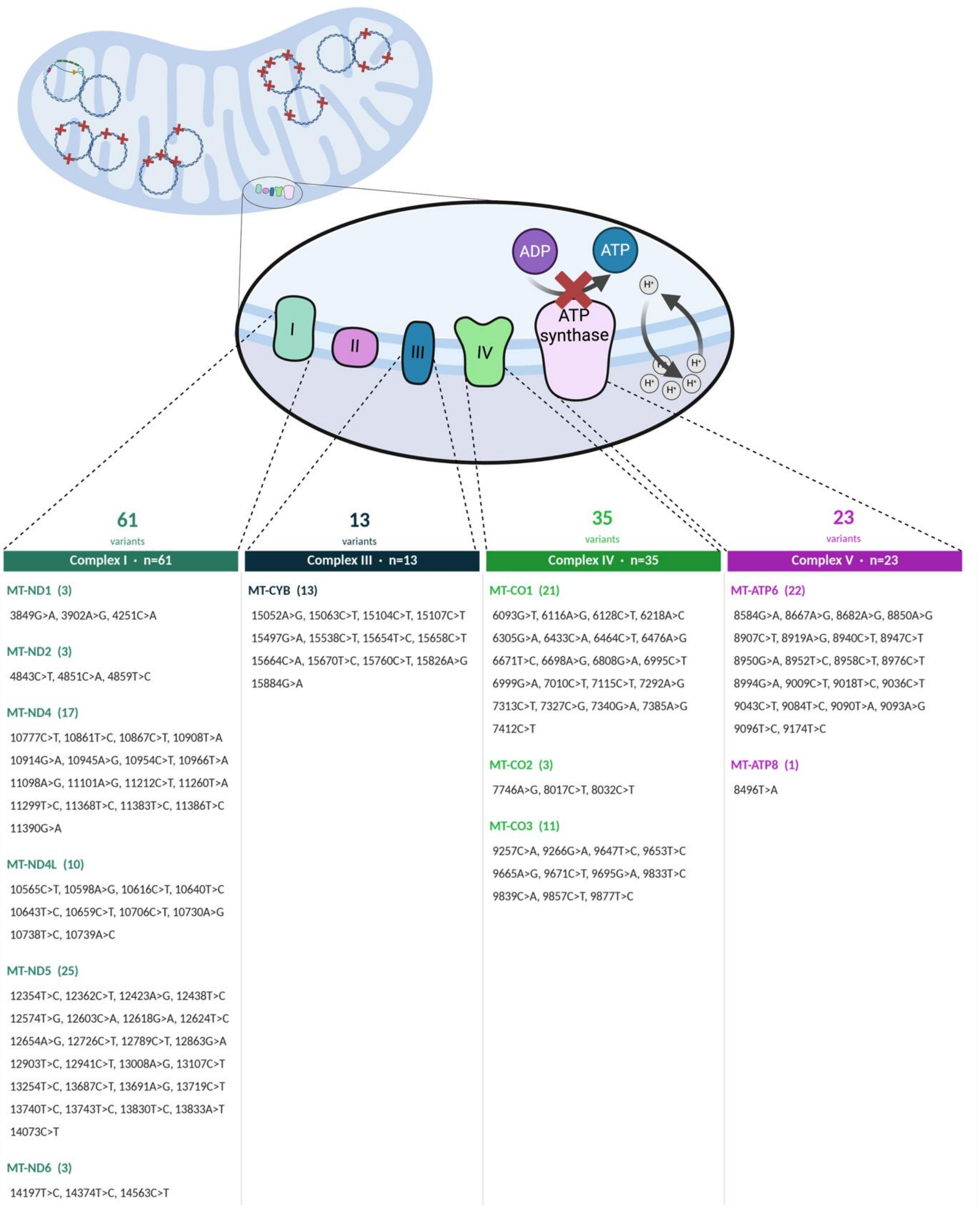


**Supplementary Figure 6. Distribution of *de novo* SNVs across mtDNA-encoded OXPHOS complexes in TOP3A KO iPSC-CMs.** Schematic of the mitochondrial OXPHOS system within the inner mitochondrial membrane, showing the distribution of *de novo* SNVs detected in TOP3A KO cells at day 90 across mtDNA-encoded subunits of complexes I, III, IV, and V. SNVs are grouped according to the OXPHOS complex indicated by the color-coded headers.

venn 1 : TOP3A KO *de novo* mtRNR variants overlapping with the cancer cohort (n = 33, 17 *MT-RNR1* + 16 *MT-RNR2*)

| Gene | Pos | Ref | Alt | Variant | De novo first detected | De novo AF D90 (%) | Cancer ShortVarID | Cancer VariantClass | Cancer Freq | Cancer Heteroplasmy |
|---|---|---|---|---|---|---|---|---|---|---|
| MT-RNR1 | 737 | C | T | 737C>T | D12 | 0.064% | C737T | 5'Flank | < 5 | 0.080 |
| MT-RNR1 | 740 | G | A | 740G>A | D12 | 0.056% | G740A | 5'Flank | 9 | 0.333 |
| MT-RNR1 | 791 | G | A | 791G>A | D12 | 0.054% | G791A | 5'Flank | < 5 | 0.373 |
| MT-RNR1 | 834 | G | A | 834G>A | D12 | 0.060% | G834A | 5'Flank | < 5 | 0.253 |
| MT-RNR1 | 889 | G | A | 889G>A | D12 | 0.054% | G889A | 5'Flank | < 5 | 0.098 |
| MT-RNR1 | 902 | G | A | 902G>A | D12 | 0.054% | G902A | 5'Flank | 7 | 0.212 |
| **MT-RNR1** | **933** | **G** | **A** | **933G>A** | **D12** | **0.051%** | **G933A** | **5'Flank** | **14** | **0.298** |
| MT-RNR1 | 954 | C | T | 954C>T | D12 | 0.090% | C954T | 5'Flank | < 5 | 0.069 |
| MT-RNR1 | 1024 | G | A | 1024G>A | D12 | 0.065% | G1024A | 5'Flank | 7 | 0.240 |
| MT-RNR1 | 1339 | G | A | 1339G>A | D12 | 0.072% | G1339A | 5'Flank | 10 | 0.432 |
| MT-RNR1 | 1344 | T | C | 1344T>C | D12 | 0.056% | T1344C | 5'Flank | < 5 | 0.331 |
| MT-RNR1 | 1389 | G | A | 1389G>A | D12 | 0.050% | G1389A | 5'Flank | 6 | 0.181 |
| MT-RNR1 | 1442 | G | A | 1442G>A | D12 | 0.048% | G1442A | 5'Flank | < 5 | 0.358 |
| MT-RNR1 | 1453 | A | G | 1453A>G | D12 | 0.045% | A1453G | 5'Flank | < 5 | 0.761 |
| MT-RNR1 | 1454 | G | A | 1454G>A | D12 | 0.060% | G1454A | 5'Flank | < 5 | 0.339 |
| MT-RNR1 | 1474 | G | A | 1474G>A | D12 | 0.059% | G1474A | 5'Flank | 6 | 0.150 |
| MT-RNR1 | 1485 | G | A | 1485G>A | D12 | 0.045% | G1485A | 5'Flank | 7 | 0.340 |
| MT-RNR2 | 1770 | G | A | 1770G>A | D12 | 0.058% | G1770A | 5'Flank | 5 | 0.243 |
| MT-RNR2 | 2008 | G | A | 2008G>A | D12 | 0.067% | G2008A | 5'Flank | 6 | 0.456 |
| MT-RNR2 | 2107 | G | A | 2107G>A | D12 | 0.067% | G2107A | 5'Flank | < 5 | 0.241 |
| MT-RNR2 | 2177 | T | C | 2177T>C | D12 | 0.065% | T2177C | 5'Flank | < 5 | 0.438 |
| MT-RNR2 | 2203 | G | A | 2203G>A | D12 | 0.064% | G2203A | 5'Flank | 5 | 0.377 |
| MT-RNR2 | 2510 | T | C | 2510T>C | D12 | 0.050% | T2510C | 5'Flank | 8 | 0.519 |
| MT-RNR2 | 2534 | G | A | 2534G>A | D12 | 0.056% | G2534A | 5'Flank | 10 | 0.268 |
| MT-RNR2 | 2560 | G | A | 2560G>A | D12 | 0.046% | G2560A | 5'Flank | < 5 | 0.236 |
| MT-RNR2 | 2592 | G | A | 2592G>A | D12 | 0.059% | G2592A | 5'Flank | < 5 | 0.242 |
| MT-RNR2 | 2759 | T | G | 2759T>G | D12 | 0.077% | T2759G | 5'Flank | < 5 | 0.064 |
| MT-RNR2 | 2935 | A | G | 2935A>G | D12 | 0.041% | A2935G | 5'Flank | < 5 | 0.266 |
| MT-RNR2 | 3032 | G | A | 3032G>A | D12 | 0.060% | G3032A | 5'Flank | 8 | 0.265 |
| MT-RNR2 | 3061 | G | C | 3061G>C | D12 | 0.083% | G3061C | 5'Flank | < 5 | 0.060 |
| MT-RNR2 | 3086 | T | G | 3086T>G | D12 | 0.063% | T3086G | 5'Flank | < 5 | 0.098 |
| MT-RNR2 | 3092 | T | C | 3092T>C | D12 | 0.047% | T3092C | 5'Flank | 5 | 0.479 |
| MT-RNR2 | 3102 | T | C | 3102T>C | D12 | 0.043% | T3102C | 5'Flank | < 5 | 0.418 |

**Supplementary Figure 7. TOP3A KO *de novo* mitochondrial rRNA SNVs overlapping the Genomics England cancer cohort.** The 33 *de novo* SNVs arising in *MT-RNR1* and *MT-RNR2* in TOP3A KO cells that were also observed in the Genomics England cancer cohort are shown (Boscenco et al, 2025). Overlap was defined by an exact match in genomic position and base substitution (reference and alternate allele). Of the 33 overlapping SNVs, 17 occur in *MT-RNR1* and 16 in *MT-RNR2*. The row highlighted in red (*MT-RNR1* m.933G>A) denotes the single overlapping SNV also reported as a recurrent SNV hotspot in the cancer cohort. All 33 SNVs were first detected at day 12.

**Supplementary Figure 8. Distribution of *de novo* and cancer-overlapping mtDNA SNVs. (A)** Total mitochondrial SNV load, comprising pre-existing and *de novo* SNVs, in TOP3A KO (red; n = 2,391) and WT (green; n = 1,209) cells at day 90 compared with *MT-RNR1*/*MT-RNR2* SNVs reported in the Genomics England pan-cancer cohort (blue; n = 1,252; Boscenco et al., 2025). Overlap required an identical genomic position and substitution, with counts separated into *MT-RNR1*, *MT-RNR2*, and other genes. **(B)** *De novo* SNVs in TOP3A KO (red; n = 304) and WT (green; n = 24) compared with the 138 recurrent SNV hotspot positions reported in the cancer cohort (blue). Four TOP3A KO *de novo* SNVs coincide with hotspot positions: *MT-RNR1* m.933G>A, *MT-RNR1* m.1336G>T, *MT-RNR2* m.3091G>T, and *MT-ND4* m.10914G>A. In the accompanying table, TOP3A KO day 90 allele frequencies are shown in red and cancer-cohort annotations in blue. At m.1336 and m.3091, the cancer cohort contains a different substitution (G>A) at the same genomic position. **(C)** Stacked-variant plot of *MT-RNR1* (12S rRNA) and adjacent *MT-TV* (tRNA-Val), m.648–1670. Red triangles denote *de novo* SNVs, purple triangles denote *de novo* SNVs also reported in the cancer cohort, and purple circles denote pre-existing SNVs also reported in the cancer cohort. Selected SNVs are labeled by position, substitution, and heteroplasmy. The plot contains 161 SNVs in total (99 pre-existing and 62 *de novo*), of which 116 positions overlap the cancer cohort, including 17 *de novo* SNVs.

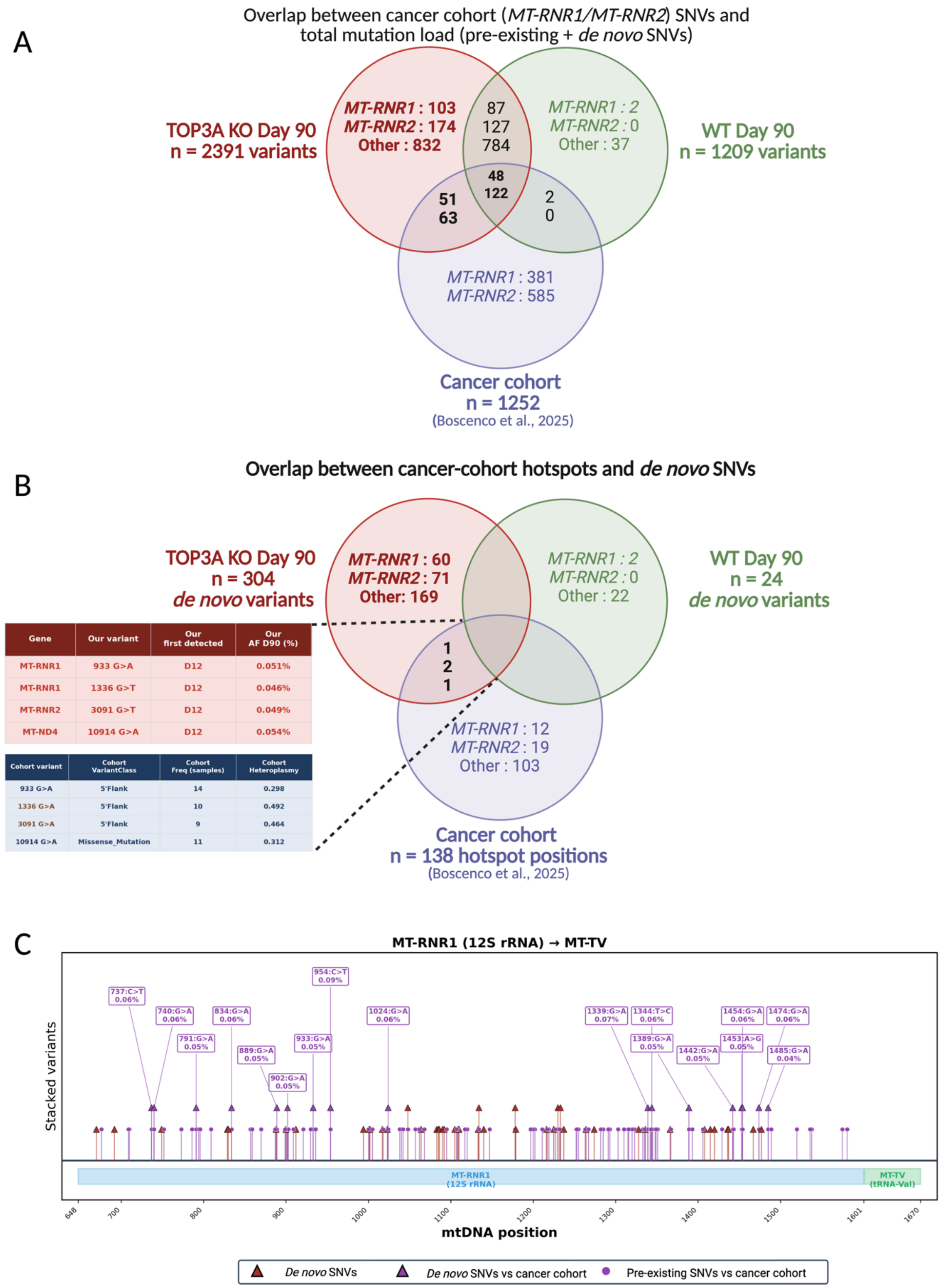

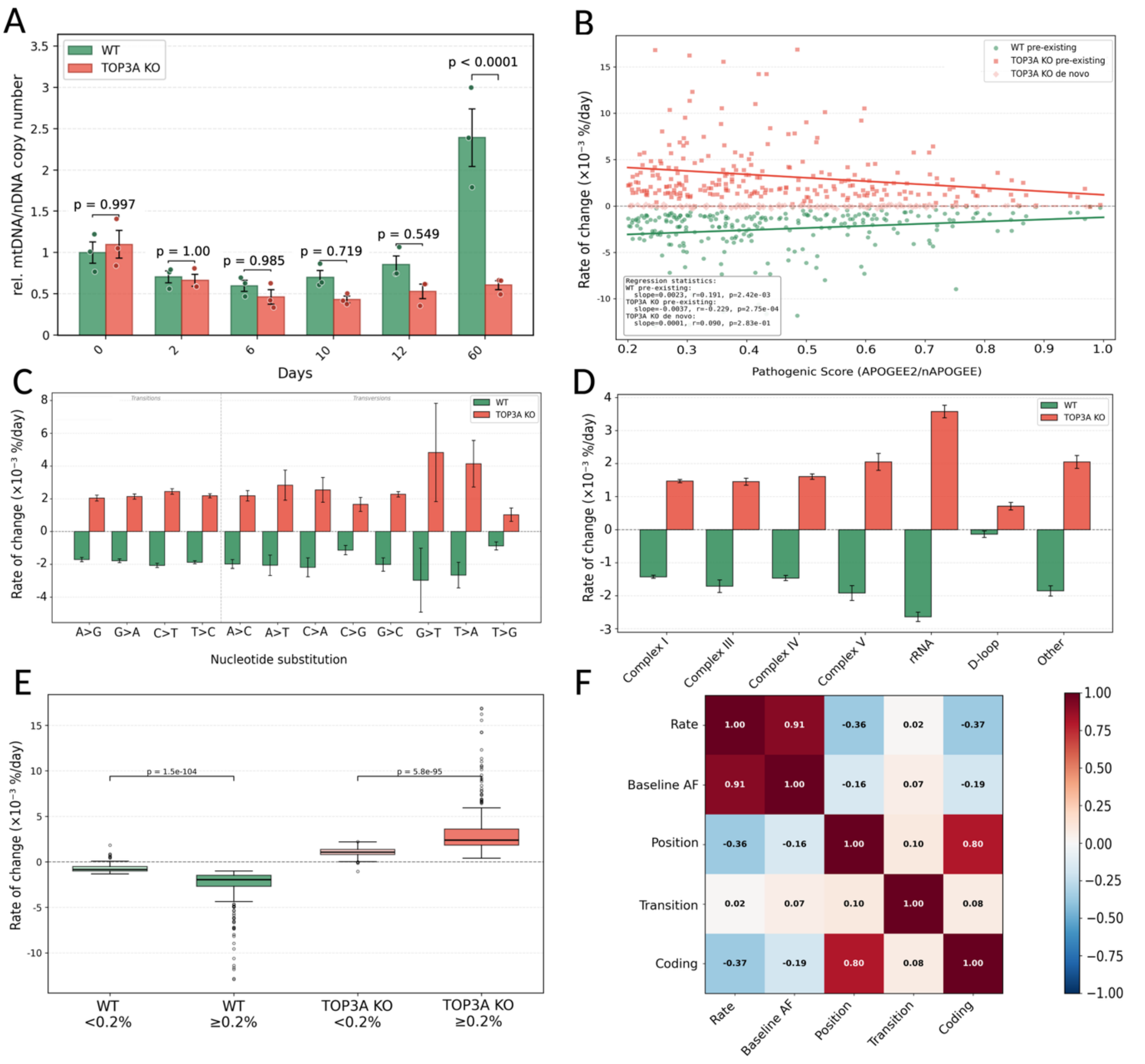


**Supplementary Figure 9. Determinants of heteroplasmy rate of change and mtDNA copy number. (A)** Relative mtDNA/nuclear DNA copy number at days 0, 2, 6, 10, 12, and 60, determined by quantitative PCR of *MT-ND1* normalized to the nuclear loci *ALB* and *F8*. Bars and error bars represent mean ± SEM, and points represent independent differentiations (n = 3 per genotype and time point). WT and TOP3A KO are shown in green and red, respectively. Genotypes were compared at each time point using two-way ANOVA followed by Šídák's multiple-comparisons test. **(B)** Heteroplasmy rate of change ($\times 10^{-3}$ %/day) versus predicted pathogenicity score for WT pre-existing variants (green), TOP3A KO pre-existing variants (red), and TOP3A KO *de novo* variants (light red). APOGEE2 scores were used for coding variants and nAPOGEE scores for noncoding variants. Lines show linear regression fits. **(C)** Mean ± SEM rate of heteroplasmy change for WT (green) and TOP3A KO (red) variants grouped by nucleotide substitution class. Transitions and transversions are separated by the vertical dashed line. **(D)** Mean ± SEM rate of heteroplasmy change for WT (green) and TOP3A KO (red) variants grouped as complex I, complex III, complex IV, complex V, rRNA, D-loop, or other. **(E)** Rate of heteroplasmy change stratified by baseline day 0 heteroplasmy below 0.2% or ≥0.2% for WT (green) and TOP3A KO (red). Boxplots show the median and interquartile range, with whiskers and outlying variant sites displayed. Within-genotype comparisons were performed using two-sided Mann–Whitney U tests. **(F)** Spearman correlation matrix of heteroplasmy rate of change, baseline allele frequency, genomic position, transition status, and coding status for shared pre-existing variants in WT and TOP3A KO (n = 723). Cells are colored according to the Spearman correlation coefficient, with the coefficient displayed within each cell.
